# Limitation of switching sensory information flow in flexible perceptual decision making

**DOI:** 10.1101/2023.12.03.569827

**Authors:** Tianlin Luo, Mengya Xu, Zhihao Zheng, Gouki Okazawa

## Abstract

Humans can flexibly change rules to categorize sensory stimuli, but their performance degrades immediately after a task switch. This switch cost is believed to reflect a limitation in cognitive control, although the bottlenecks responsible for this remain controversial. Here, we show that humans exhibit a brief reduction in the efficiency of converting sensory inputs into decision evidence immediately after changing rules in perceptual tasks. Participants performed a flexible face categorization task in which they classified parametrically generated face stimuli based on one of two rules, switching every few trials. Although participants were always informed of a rule switch by a context cue, they showed a specific pattern of increase in reaction times, regardless of the time they were given to prepare for the switch. Psychophysical reverse correlation and computational modeling revealed a reduction in sensory weighting immediately after a rule switch, which recovered within a few hundred milliseconds after stimulus presentation. Furthermore, this cost depends on the sensory features being switched, suggesting a major bottleneck at the stage of adjusting the sensory information flow. We propose that decision-making circuits cannot fully adjust their sensory readout based on an abstract context cue alone, but rather require the presence of an actual stimulus to tune it, leading to a limitation in flexible perceptual decision making.

## Introduction

Successful sensory-guided behavior relies on the ability to transform relevant sensory signals into appropriate action plans that meet task demands. Decades of work have investigated this behavior using perceptual decision-making tasks with fixed stimulus-action mapping, such as the direction discrimination of stochastic moving dots (Shadlen and Kiani 2013). Under these conditions, many aspects of behavior, such as choices, reaction times (RTs), and confidence associated with choices, can be accurately explained by a class of models that accumulate sensory evidence toward a decision bound (Forstmann et al. 2016; Gold and Shadlen 2007; Ratcliff et al. 2016; Shadlen and Kiani 2013). These models have been further supported by the findings of neural activity in multiple brain regions that reflect similar computational processes (Gold and Shadlen 2007; Hanks and Summerfield 2017; O’Connell and Kelly 2021).

An important problem that has not been addressed in these frameworks is how the brain flexibly selects relevant sensory information depending on the behavioral context (Okazawa and Kiani 2023). Our tasks in the real world are diverse and variable, and our brains must constantly adjust the association of sensory inputs and actions. Recent neurophysiological studies have begun to investigate this selection process by employing task designs that require subjects to switch between two perceptual tasks, such as color versus motion discrimination of colored random dot motions (Mante et al. 2013; Sasaki et al. 2022; Siegel et al. 2015; Suda and Uka 2022; Tsumura et al. 2021). These studies have found that neural population signals reflect flexible sensory gating (Flesch et al. 2022; Mante et al. 2013; Pagan et al. 2022; Soldado Magraner et al. 2023). Simultaneously, modeling frameworks using recurrent neural networks (RNNs) have been developed to account for neural activity (Mante et al. 2013; Yang et al. 2019). RNNs often instantiate selection mechanisms through internal dynamics that switch the flow of sensory information according to external task context signals.

However, humans exhibit notable behavioral limitations in switching tasks that are absent in these network models. Immediately after a task switch, decisions often become less accurate and slower (Karayanidis et al. 2010; Kiesel et al. 2010; Koch et al. 2010; Monsell 2003), even when the switch is predictable (Rogers and Monsell 1995) or explicitly cued (Meiran 1996). This switch cost is considered an important property of the brain’s mechanisms of cognitive control (Flesch et al. 2023; Musslick and Cohen 2021). A common explanation is that the brain needs time to reconfigure its internal state for a switched rule (Ardid and Wang 2013; Jaffe et al. 2023; Monsell 2003; Rogers and Monsell 1995) or to suppress the effect of the previous task rule arising from inertia (Allport et al. 1994) or priming (Mayr and Kliegl 2003) from the previous internal state. If preparation time is insufficient, this control process interferes with the subsequent decision-making process and affects task performance (Monsell 2003). Interestingly, even when sufficient time is given after a task switch is cued, humans still exhibit substantial switch costs (residual switch cost; de Jong 2000; Li et al. 2019; Meiran et al. 2000; Monsell and Mizon 2006; Rogers and Monsell 1995), suggesting that the brain is unable to fully adjust its internal state based on external cues alone. This has also been explained as a limitation in cognitive control: either participants fail to engage in a task until the task begins (de Jong 2000), they cannot fully switch attention (Elchlepp et al. 2017, 2015; Longman et al. 2014), or the task stimuli themselves are required to recall the stimulus-response mapping (Koch and Allport 2006; Logan and Bundesen 2004; Schneider and Logan 2005). However, a satisfactory explanation for why a contextual cue is insufficient for the brain to switch its state internally is still lacking. This fundamental constraint on task switching provides an important clue regarding how neural circuits implement computational flexibility.

Here, we show that such switch costs in perceptual decision making reflect a reduction in the efficiency of converting sensory inputs to decision evidence immediately after a task switch. This efficiency reduction cannot be ameliorated by longer task preparations, but quickly recovers to the baseline level within a few hundred milliseconds after stimulus presentation. Furthermore, the magnitude of the switch cost substantially depends on the specific sensory features to be switched, discordant with the idea that the cost is primarily caused by the transition of abstract cognitive states. These findings were obtained by employing advanced behavioral measurement and modeling techniques to study perceptual decision making (Fetsch 2016; Okazawa et al. 2021b, 2018; Waskom et al. 2019). In this task, participants switched categorization rules for parametrically morphed facial stimuli that had stochastic evidence fluctuations during stimulus presentation. Psychophysical reverse correlation and computational modeling revealed an initial reduction in sensory weighting that resulted in a switch cost. We suggest that when switching relevant sensory dimensions, decision-making circuits cannot fully adjust their sensory readout based on an abstract context cue alone, but require the presence of an actual stimulus to fine-tune the readout for certain sensory features.

## Results

### Switch cost independent of stimulus strength and task preparation time

We developed a context-dependent face categorization task, in which participants classified a face stimulus based on one of two task rules. We used face categorization because previous studies successfully explained the behavior using a simple evidence accumulation model (Okazawa et al. 2021b, 2018) and the high-dimensional nature of face stimuli allowed us to easily introduce flexibility in the task, such as switching between identity and expression categorization (Berger et al. 2019; Elchlepp et al. 2021; Schuch et al. 2012).

In each trial, participants first fixated on a central fixation point whose color indicated the task rule, then viewed a face stimulus sampled from a two-dimensional (2D) morphed face space (Figs. 1A-B), and reported the face category by making a saccade to one of the two targets as soon as they were ready. The two rules correspond to the two axes of 2D space (e.g., identity vs. expression; Fig. 1A). The category boundary was at the center of each axis (0% morph level) and the stimulus became easier as the distance from the boundary (absolute value of the morph level) increased. Importantly, on each trial, the morph levels of the face stimulus fluctuated randomly every 106.7 ms around the sampled point in the 2D space (Fig. 1B inset), allowing us to estimate how participants temporally weighted each stimulus frame to make a decision (i.e., psychophysical reverse correlation; Ahumada Jr 1996). Each face frame transition was interleaved with a mask image so that the fluctuations remained subliminal. This task design allowed us to quantitatively compare differences in decision-making processes between the trials immediately following a rule switch (“switch” trials) and the remaining “non-switch” trials (Fig. 1C; task switched every 2-6 trials following truncated exponential distribution).

**Figure 1:**
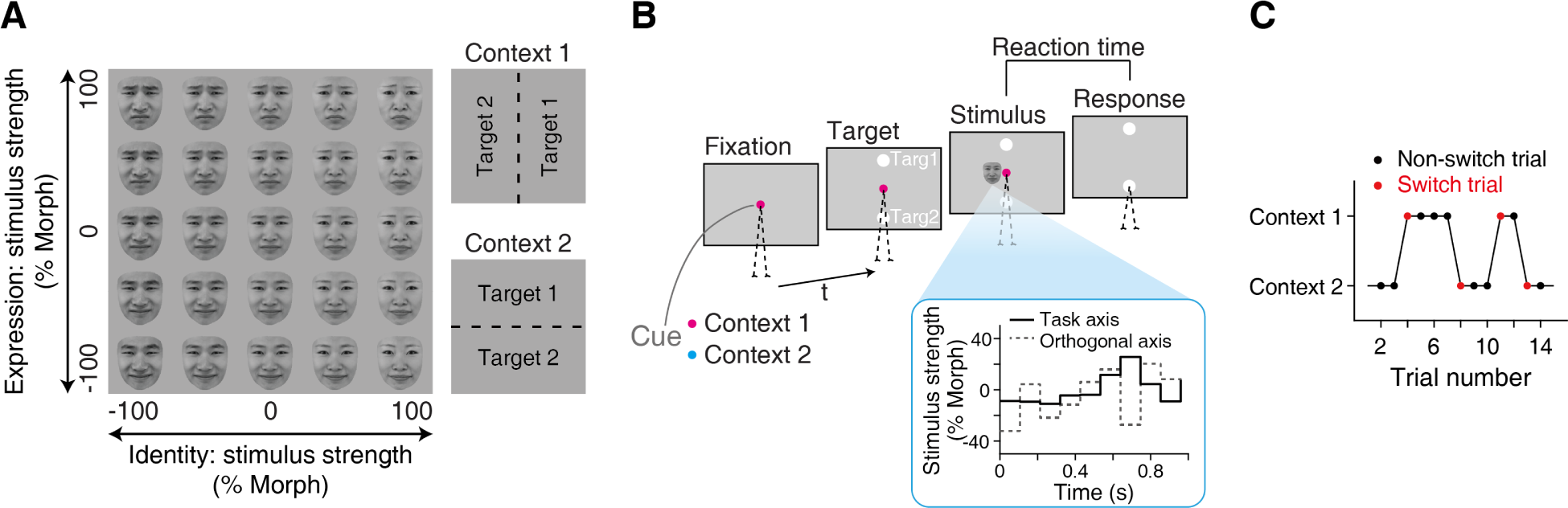
Context-dependent face categorization task. (**A**) A 2D face stimulus space. Each axis corresponds to one of the two categorization rules (e.g., identity vs. expression). In each rule, a category boundary divides the stimulus space into two halves, which were associated with two saccade targets (right column). Six out of the eight participants performed identity vs. expression categorization, while the remaining two performed identity vs. age categorization (Supplementary Fig. 1). The prototype faces were from the Tsinghua Facial Expression Database (Yang et al. 2020) and presented with permission. (**B**) Participants initiated each trial by fixating on a central point whose color indicated the current task rule. Shortly after, two target points appeared, followed by a sequence of face stimuli. In the sequence, the morph levels of face stimuli were randomly updated every 106.7 ms—drawn from a Gaussian distribution with a mean chosen for each trial and SD 20% morph level along both task-relevant and orthogonal axes—providing noisy sensory evidence (inset). Participants reported the stimulus category by making a saccade to one of the two targets as soon as they were ready. Reaction time was defined as the time between the stimulus onset and the saccade onset. (**C**) Two task contexts were switched every 2-6 trials (truncated exponential distribution). The trials immediately following a rule switch were defined as switch trials (red), while the remaining trials were classified as non-switch trials (black).

After sufficient training, participants could switch task rules immediately without affecting their choice accuracy, but their RTs were substantially longer during the switch trials. Psychometric functions along the task-relevant axis were nearly indistinguishable between switch and non-switch trials (Fig. 2A left; *α*_1_*_s_* = *−*1.1 *±* 0.5 in Eq. 2, mean *±* S.E.M. across participants; *t*_(7)_ = *−*2.3, *p* = 0.053, two-tailed *t*-test) and had negligible lapse rates for the easiest stimuli (*<* 1.2% for all participants), suggesting that participants almost never confused the task rule. Mean RTs were faster for easier stimuli as in typical perceptual tasks (Fig. 2A right; *β*_2_ = 5.2 *±* 0.4 in Eq. 3; *t*_(7)_ = 14.3, *p* = 9.6 *×* 10*^−^*^7^, right-tailed *t*-test), but they were systematically longer in switch trials (Fig. 2A; *β*_0_*_s_* = 0.17 *±* 0.02; *t*_(7)_ = 8.7, *p* = 2.7 *×* 10*^−^*^5^; *∼*170 ms longer on average across stimulus strengths).

**Figure 2:**
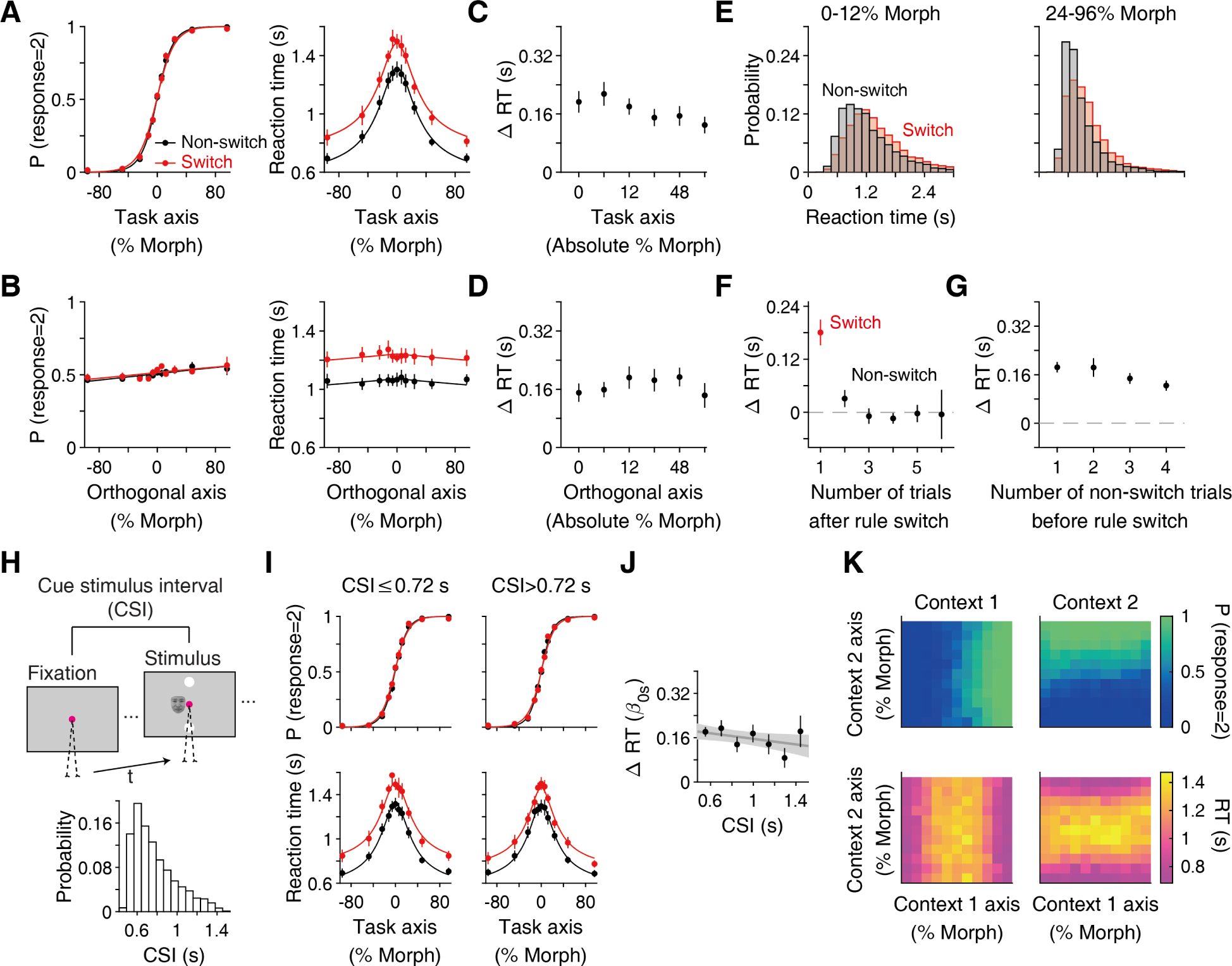
Task switching prolonged reaction times regardless of stimulus difficulty and preparation time. (**A**) Reaction times (RTs) increased for all stimulus strengths in the switch trials (right), while choice accuracy was nearly identical between the switch and non-switch trials (left). Smooth lines are fit by a logistic function (left; Eq. 2) and a hyperbolic tangent function (right; Eq. 3). The plots are the average of the eight participants. (**B**) Choice accuracy and RTs did not depend on the stimulus strength along the task-orthogonal axis. (**C, D**) The RT increase was independent of the stimulus strengths both along the task axis (C) and orthogonal axis (D). (**E**) For difficult stimuli (left), the RT distribution shifted positively from non-switch to switch trials, while for easy stimuli (right), the RT distribution flattened. (**F, G**) The increase in RTs was present only in the first trial after a task switch (F) and did not depend on the number of non-switch trials before the switch (G). The plots are the average of all the stimulus strengths. (**H**) Participants could prepare for the switched rule after fixating on a context cue until stimulus onset. This cue-stimulus interval (CSI) approximately followed a truncated exponential distribution in our task (bottom). (**I**) The increase in RTs after a task switch occurred both for the trials with short (left) and long (right) CSIs. Trials were spilt at the median CSI (0.72 s). (**J**) The increase in RTs (defined as β_0s_ in Eq. 3) was independent of CSIs. The line is a linear regression averaged across participants. (**K**) Psychometric (top) and chronometric (bottom) functions in the 2D space revealed no congruency effect in our task. The stimuli in the second and fourth quadrants of the face space were incongruent, as they were associated with the opposite targets between the two tasks. Participants showed neither lower accuracy nor longer RTs for these stimuli.

Notably, the increase in average RTs was largely independent of the stimulus difficulty. We computed the difference in average RTs between the switch and non-switch trials and confirmed that it was minimally correlated with stimulus difficulty (Fig. 2C; *F*_(5_,_42)_ = 1.4, *p* = 0.25, one-way ANOVA, the main effect of morph level along the task axis). The increase in RTs occurred regardless of the direction of task switching (e.g., identity to expression or expression to identity; see Fig. 6F) but was limited to only one trial following a switch, and no obvious effect was observed thereafter (Fig. 2F; *p >* 0.15 for all subsequent trials before the next task switch, two-tailed *t*-test). Furthermore, the RT increase did not depend on the number of non-switch trials preceding a switch (Fig. 2G; *F*_(3_,_28)_ = 1.9, *p* = 0.16, one-way ANOVA). Thus, participants correctly switched the task rule and maintained choice accuracy, but their average RTs increased uniformly across a range of stimulus difficulties only at the moment of the task switch. However, when we plotted the RT distributions for difficult and easy stimuli, it appeared as a shift in the distribution for difficult stimuli, but a flattening of the distribution for easier stimuli (Fig. 2E), implying that the RT increase was not merely an addition of a fixed amount of time.

Indeed, we found evidence that this increase in average RTs cannot be explained by the additional time required to prepare for switching rules. Importantly, the RT increase occurred regardless of the preparation time allotted to participants for a task switch. In our task, the fixation point color indicated the task rule to the participants, and a stimulus was presented after a variable duration following the participant’s fixation onset (cue-stimulus interval, CSI: range, 0.5-1.5 s; median, 0.72 s; truncated exponential distribution; Fig. 2H). If the CSI is not long enough for participants to prepare for a task switch, shorter CSIs should result in less task preparation and a delay in initiating the decision-making process. Inconsistent with this prediction, we found that CSIs had no significant impact on the increase in RTs within the range of tested CSIs (Figs. 2I-J; slope = *−*0.05 *±* 0.05; *t*_(7)_ = *−*1.1, *p* = 0.33, two-tailed *t*-test). In other words, participants consistently spent an additional *∼*170 ms after stimulus onset to make a decision in the switch trials, despite the much longer and variable preparation time available before stimulus onset.

Previously, this persistent behavioral effect of task switching has been termed the residual switch cost (de Jong 2000; Meiran et al. 2000; Monsell and Mizon 2006; Rogers and Monsell 1995). If CSIs are shorter than the range we used (*<*0.5 s; Meiran et al. 2000), the switch cost can become more substantial, which likely reflects the time participants needed to prepare for a new task (task set reconfiguration; Monsell 2003) or suppress previous rules (Allport et al. 1994; Yeung and Monsell 2003). However, it remains controversial as to why the cost persists as a residual switch cost with a longer preparation time (de Jong 2000; Elchlepp et al. 2021; Schneider 2016).

One hypothesis is that the stimulus itself triggers the reconfiguration of task rules (Koch and Allport 2006; Logan and Bundesen 2004; Schneider and Logan 2005). However, the stimuli in our task contained no clues to task rules, as they were sampled from the same 2D face space in both contexts. If participants perceived stronger sensory signals along the task-relevant axis as a task cue (e.g., clearly happy or sad faces triggered the use of the expression rule), higher morph levels would lead to smaller switch cost, but as shown above, the increase in RTs was independent of stimulus strength (Figs. 2A, C). The increase in RTs was also not correlated with the morph levels of the task-orthogonal axis (Figs. 2B, D; *F*_(5_,_42)_ = 0.6, *p* = 0.68, one-way ANOVA, the main effect of stimulus strength along the orthogonal axis), ruling out the possibility that stronger task-orthogonal inputs caused task confusion. Furthermore, RTs were not particularly long for stimuli associated with the opposite saccade targets in the two contexts (i.e., incongruent stimuli) (Fig. 2K; congruent stimuli, RT = 1.07 *±* 0.04 s; incongruent stimuli, RT = 1.06 *±* 0.04 s; *t*_(7)_ = *−*0.48, *p* = 0.68, right-tailed paired *t*-test), indicating that RT increases were not due to conflict at the response level.

Then, why does task switching prolong RTs regardless of stimulus strengths without affecting choice accuracy? Previous studies established a simple evidence accumulation model that accurately accounted for choices and RTs during face categorization (Okazawa et al. 2021b, 2018). Such a modeling approach is suitable for examining the mechanistic components that explain the observed behavioral changes after task switching. Furthermore, random stimulus fluctuations in our task (Fig. 1B) enabled us to test whether and how participants changed the weighting of sensory evidence for their decisions in the switch trials. In the next section, we demonstrate that RTs increase owing to a brief initial reduction in sensory weighting.

### Brief initial reduction in sensory processing efficiency explains switch cost

We performed a psychophysical reverse correlation (Ahumada Jr 1996; Okazawa et al. 2018) to examine how temporal stimulus fluctuations influenced participants’ behaviors in the switch and non-switch trials. In brief, we calculated the difference in average morph fluctuations between trials in which participants chose one target over another (Eq. 4). The amplitudes of the resulting psychophysical kernels reflect the degree to which sensory fluctuations at each moment influenced the participants’ choices; thus, they are informative for estimating how participants weigh sensory evidence to make a decision (Okazawa et al. 2018). The kernels were aligned to either the stimulus onset or the timing of the participants’ saccadic responses (Fig. 3A).

**Figure 3:**
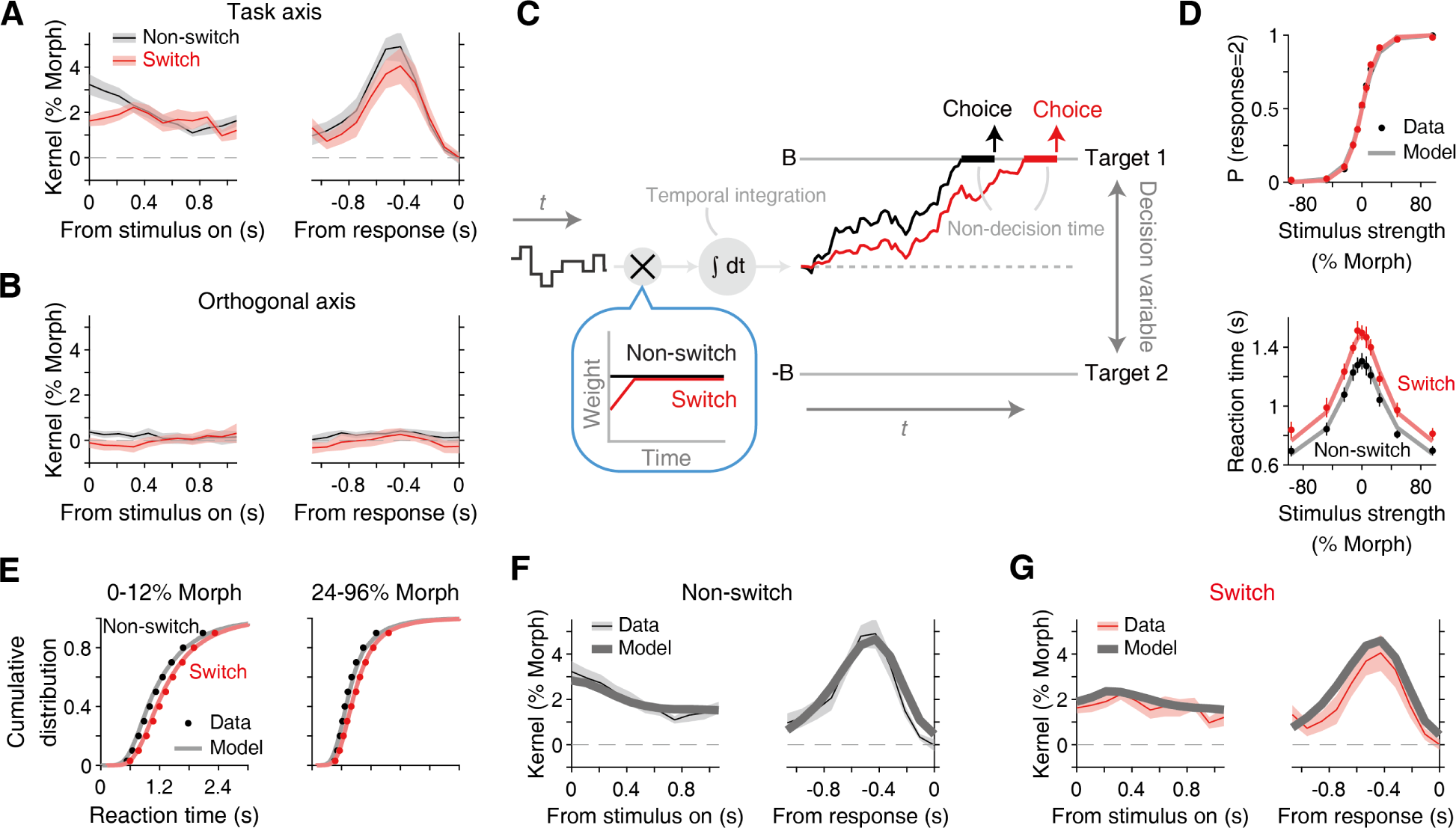
A brief initial reduction in sensory weight accounts for choices, reaction times, and psychophysical kernels. (**A**) Psychophysical kernels (Eq. 4) along the task axis aligned to stimulus onset (left) and participants’ response (right). The dynamics of the kernels for the non-switch trials (black) were similar to those observed previously and consistent with a bounded evidence accumulation mechanism (Okazawa et al. 2018). The kernels for the switch trials (red), by comparison, showed a brief reduction in amplitude at stimulus onset. Shading indicates S.E.M. across participants. (**B**) The kernels along the task-orthogonal axis were indistinguishable from zero, indicating that the orthogonal information did not influence participants’ decisions in both switch and non-switch trials. (**C**) An evidence accumulation model that accounts for both switch and non-switch trials. The model receives fluctuating sensory information, integrates the evidence toward decision bounds and, when it reaches a bound, commits to the choice associated with that bound. Reaction time is the sum of bound crossing time and non-decision time, including sensory and motor delays. In switch trials, we added a brief reduction in sensory weight (inset). (**D, E**) The model accurately fits choices (D, top), mean reaction times (D, bottom), and reaction time distributions (E; cumulative distributions are shown for visualization) for both switch and non-switch trials. (**F, G**) The same model accurately explains the patterns of the psychophysical kernels. The data traces are the same as that in A and B.

In the non-switch trials, we found dynamic kernel patterns consistent with linear evidence accumulation. The kernel aligned to stimulus onset showed a gradual decrease in amplitude, whereas the kernel aligned to the participants’ saccade revealed a characteristic peak a few hundred milliseconds before the saccade (black line in Fig. 3A). Although these patterns seem to imply the dynamic weighting of sensory evidence, previous studies have shown that they can be fully explained by a bounded evidence accumulation mechanism with a constant sensory weight (Okazawa et al. 2021b, 2018). In this model (Fig. 3C), sensory fluctuations are linearly accumulated toward the upper or lower bound. When the accumulated evidence reaches a bound, this bound determines the choice. RTs are modeled as the time required to accumulate evidence plus the time irrelevant to decision making (non-decision time) including sensory and motor delays.

This evidence accumulation model accurately fit the psychometric (Fig. 3D top; *R*^2^ = 0.98 *±* 0.005) and chronometric functions (Fig. 3D bottom; *R*^2^ = 0.95 *±* 0.007) of the non-switch trials as well as the RT distributions (Fig. 3E). The same model also quantitatively explained the psychophysical kernels. Using the fitted model parameters, we simulated the model responses to randomly generated stimulus fluctuations and computed the psychophysical kernels of the model (thick gray line in Fig. 3F; *R*^2^ = 0.91 with the actual data). This model explains the gradual reduction of the kernel aligned to the stimulus onset (Fig. 3F, left) because there is a temporal gap between the bound crossing and the report of a decision (i.e., the non-decision time). This non-decision time renders a later portion of the stimulus fluctuations irrelevant to the decision. Because the timing of the bound crossing varies across trials, the model predicts a gradual reduction in the effect of stimulus fluctuations over time (Okazawa et al. 2018). The model also explained the peak of the kernel aligned with the participant’s saccade (Fig. 3F, right). This peak arises because, near the time of bound crossing, tiny stimulus fluctuations can push the decision variable beyond the bound and dictate the decision. Therefore, at that moment, the effect of stimulus fluctuations becomes substantial and peaks. After this peak, the kernel drops sharply to zero because of the non-decision time (Okazawa et al. 2018; Zylberberg et al. 2012).

Now, in the switch trials, we found that the kernel aligned to the stimulus onset showed a characteristic reduction in amplitude during the first 200–300 ms after stimulus onset compared to that of the non-switch trials (red line in Fig. 3A; *t*_(7)_ = 3.3, *p* = 0.0065, right-tailed paired *t*-test at the first stimulus frame). The amplitude reduction was approximately 39% (Eq. 5) from the non-switch trials, and then recovered over time. Similar to the increase in RTs, this kernel reduction was independent of the cue-stimulus interval (CSI *≤* 0.72 vs. *>* 0.72, *t*_(7)_ = *−*0.54, *p* = 0.61, two-tailed paired *t*-test), and thus occurs regardless of the preparation time. Aside from this initial reduction, there was no noticeable difference in the kernels between switch and non-switch trials.

Inspired by the observed kernel patterns, we added a dynamic sensory weighting function to the evidence accumulation model (Fig. 3C inset; Eq. 11). The weight was constant in the non-switch trials; however, in the switch trials, it is a ramp function that starts with an initially reduced amplitude *w*_0_ and recovers to the baseline level at time *t_w_*. While this function modulates sensory evidence, our model is agnostic of whether such modulation occurs during sensory processing or during the conversion of sensory information into accumulated evidence (see Discussion section). Hereafter, we collectively refer to this as a reduction in the efficiency of processing sensory evidence. To test if this efficiency reduction alone was sufficient to account for the behavioral changes in the switch trials, we started with the model parameters fitted to the non-switch trials and allowed the model to change only these two additional parameters (*w*_0_ and *t_w_*) to fit the behavioral data in the switch trials.

This extended model accurately accounted for participants’ choices (Fig. 3D top; *R*^2^ = 0.98 *±* 0.006), mean RTs (Fig. 3D bottom; *R*^2^ = 0.88 *±* 0.031), and RT distributions in the switch trials (Fig. 3E). The fitted parameters showed approximately half the amplitude of sensory sensitivity at the stimulus onset on switch trials (*w*_0_ = 0.42 *±* 0.21) that recovered in a few hundred milliseconds (full recovery time: *t_w_* = 637 *±* 237 ms). This led to a reduction in the amplitude of the simulated psychophysical kernels, which was in good agreement with the observed data (Fig. 3G; *R*^2^ = 0.69). This reduced sensory weight delayed the time required to reach a bound, resulting in longer average RTs. The RT distribution shifted horizontally for difficult stimuli and broadened for easy stimuli, which was successfully replicated by the model (Fig. 3E). Note that these model outcomes also depended on how the weighting function modulated noise in the accumulation process (Eq. 12 and 13), as discussed in Supplementary Figure 2. In contrast to RTs, choice accuracy was almost unaffected because the reduction in sensory weight was transient and sufficient evidence could be accumulated during the subsequent long integration time. Overall, this simple addition to the decision-making model quantitatively accounted for behavioral patterns in switch trials.

We further confirmed that no other mechanisms accounted for the observed behavioral results. Multiple parameters in the evidence accumulation model can increase RTs; however, changing these parameters yields choice accuracy, RTs, and psychophysical kernels that are distinct from the observed data (Fig. 4 and Supplementary Fig. 3). For example, increasing the non-decision time in the model (Fig. 4A) prolongs RTs uniformly across stimulus strengths without affecting choice accuracy, thus explaining the observed choice and mean RTs (Fig. 4B) but failing to account for the patterns of the psychophysical kernels (Fig. 4C). A longer non-decision time does not produce an initial reduction in the onset-aligned kernel but shifts the peak of the response-aligned kernel that reflects the timing of the bound crossing. However, this pattern was not observed in the actual data.

**Figure 4:**
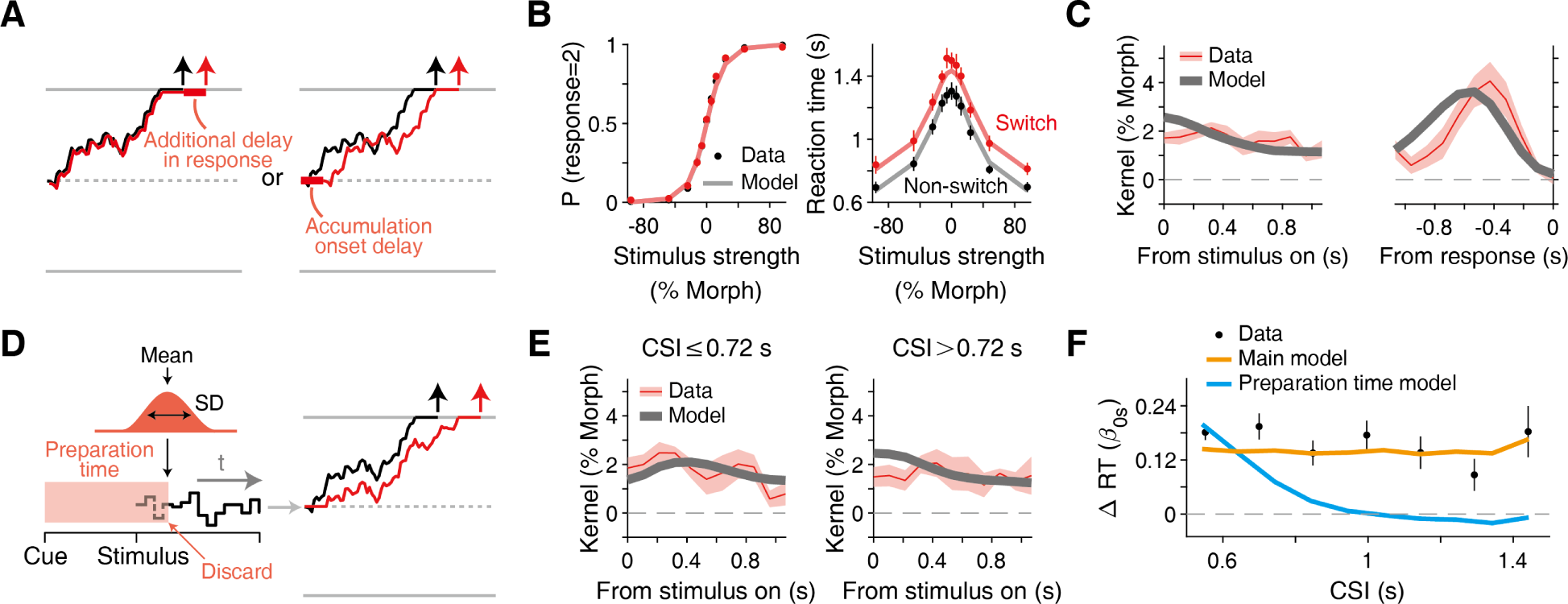
Other models fail to explain the behavioral data. (**A**) A model that explains switch cost by increasing non-decision time. This could happen either due to a longer duration of executing action after committing to a choice (left) or due to a longer delay before initiating the evidence accumulation (right). In contrast to our main model (Fig. 3C), rule switch does not affect the accumulation time. (**B, C**) The model successfully explains the lack of change in accuracy (B, left) and the prolonged reaction times irrelevant to stimulus strength (B, right), but fails to explain the patterns of psychophysical kernels (C). A longer non-decision time shifts the peak of the response-aligned kernel, while it does not lead to the initial drop of the kernel aligned to the stimulus onset. (**D**) A model that explains switch cost based on task preparation. It assumes that a fixed amount of time (following a Gaussian distribution) is required to prepare for a task switch after cue onset. If this preparation time takes longer than a cue-stimulus interval (CSI), it overlaps an initial part of the stimulus sequence, which is ignored in the decision-making process. (**E**) The task preparation model fails to fit the initial reduction of the stimulus-aligned kernel for long CSIs (CSI>0.72 s, right panel). (**F**) The model also fails to explain that the increase in reaction times is independent of CSIs. The black dots are the data (same as those in Fig. 2J). The blue line is the fit of the task preparation model averaged across participants. The orange line is the fit of the main model (Fig. 3C).

Another alternative is a leak in the evidence accumulation process (Supplementary Fig. 3B), which leads to a reduced amplitude of the kernel near the onset of the stimulus (Okazawa et al. 2018). However, this results in an increase in RTs for difficult stimuli because they require more time to reach a bound and are thus more affected by leakage. This change in RTs is inconsistent with the observed data. Similarly, increasing the decision bound or decreasing the sensory sensitivity (drift rate) (Supplementary Figs. 3C-E) led to longer RTs, but their patterns and effects on choice accuracy and psychophysical kernels were distinct from the data. In brief, higher decision bounds improve the overall accuracy and increase RTs, especially for more difficult stimuli. Lower sensory sensitivities deteriorate overall accuracy and increase RTs, especially for easier stimuli. A combination of the two evenly increases RTs for all difficulty levels with little effect on choice accuracy. However, they reduce the overall amplitude of the psychophysical kernels (Okazawa et al. 2018) instead of only reducing the initial part of the kernel. Thus, none of these alternative model parameters satisfactorily account for the effects of task switching.

Our modeling framework could also quantitatively confirm that the observed switch cost is not owing to the lack of sufficient task preparation time prior to the stimulus presentation (Figs. 4D-F). Suppose that participants need time to prepare for a task switch based on a context cue, and if the preparation time exceeds a cue-stimulus interval (CSI), they cannot initiate decision formation and ignore the initial part of the stimulus sequence (Fig. 4D). We modeled the preparation time as a Gaussian distribution and fitted the switch trials. This model showed poorer fitting performance (the difference in log-likelihood from the main model: *−*27 *±* 10, corresponding to *∼* 1.9 *×* 10*^−^*^12^ less likely; *n* = 8, *p* = 0.0039, left-tailed Wilcoxon signed-rank test). As expected, the model predicted a smaller reduction in the initial portion of the stimulus-aligned kernel with longer CSIs, which was inconsistent with the data (Fig. 4E). Accordingly, it systematically deviated from the data, which showed the independence of CSIs and the increase in RTs (Fig. 4F).

Finally, we found no evidence of interference from the task-irrelevant rule. Because our stimuli had the same degree of stimulus fluctuations along the task-orthogonal axis, we could perform a psychophysical reverse correlation using these orthogonal fluctuations. The resulting kernels had nearly zero amplitude throughout the stimulus presentation period (Fig. 3B). This suggests that the initial weight reduction was not due to residual attention or faulty accumulation of task-orthogonal information.

Altogether, we showed that longer RTs in the switch trials occurred because of the reduced efficiency of processing sensory evidence at stimulus onset, which recovered within a few hundred milliseconds. This reduction was evident in psychophysical kernels, and our modeling framework confirmed that this mechanism alone was sufficient to explain all aspects of behavioral changes from non-switch to switch trials.

### Task switching impairs choice accuracy when stimulus duration is limited

A key implication of the above observations is that task switching influences evidence accumulation in perceptual decision making. However, it did not affect choice accuracy, as opposed to findings from multiple previous studies (Rogers and Monsell 1995; Tsumura et al. 2021), because participants were able to continue accumulating evidence after the sensory weight recovered, thus maintaining performance at the expense of longer RTs (Fig. 3C). This interpretation predicts that when the stimulus duration is externally constrained by the environment, participants should now show impaired accuracy in switch trials, whereas the accuracy would not change if the increased RTs were due to a process irrelevant to decision formation, such as motor preparation.

To test this prediction, we conducted a modified experiment (Fig. 5A) in which we fixed the stimulus duration to 320-640 ms (in steps of 106.7 ms stimulus frames, following a geometric distribution), while keeping the other experimental parameters identical. In line with our prediction, we observed reduced choice accuracy in the switch trials (Fig. 5B; *α*_1_*_s_* = *−*2.2 *±* 0.8 in 6; *t*_(3)_ = *−*2.6, *p* = 0.039, left-tailed *t*-test). The psychophysical thresholds were systematically higher in switch trials for a range of the CSIs (Fig. 5C; *F*_(1_,_24)_ = 16.9, *p* = 4.0 *×* 10*^−^*^4^, repeated-measure two-way ANOVA, the main effect of switch vs. non-switch), and this was independent of the CSIs (*F*_(3_,_24)_ = 0.2, *p* = 0.87, the main effect of CSIs). Furthermore, we still observed a small increase in RTs in the switch trials (Fig. 5D; *β*_0_*_s_* = 0.04 *±* 0.01; *t*_(3)_ = 4.5, *p* = 0.010, right-tailed *t*-test; *∼*38 ms longer on average across stimulus strengths). This was expected from the model because the probability of reaching a decision bound before stimulus termination is lower in switch trials (Kiani et al. 2008). Thus, task switching affects decision formation, leading to longer RTs or lower accuracy, depending on the accessibility to further sensory inputs.

**Figure 5:**
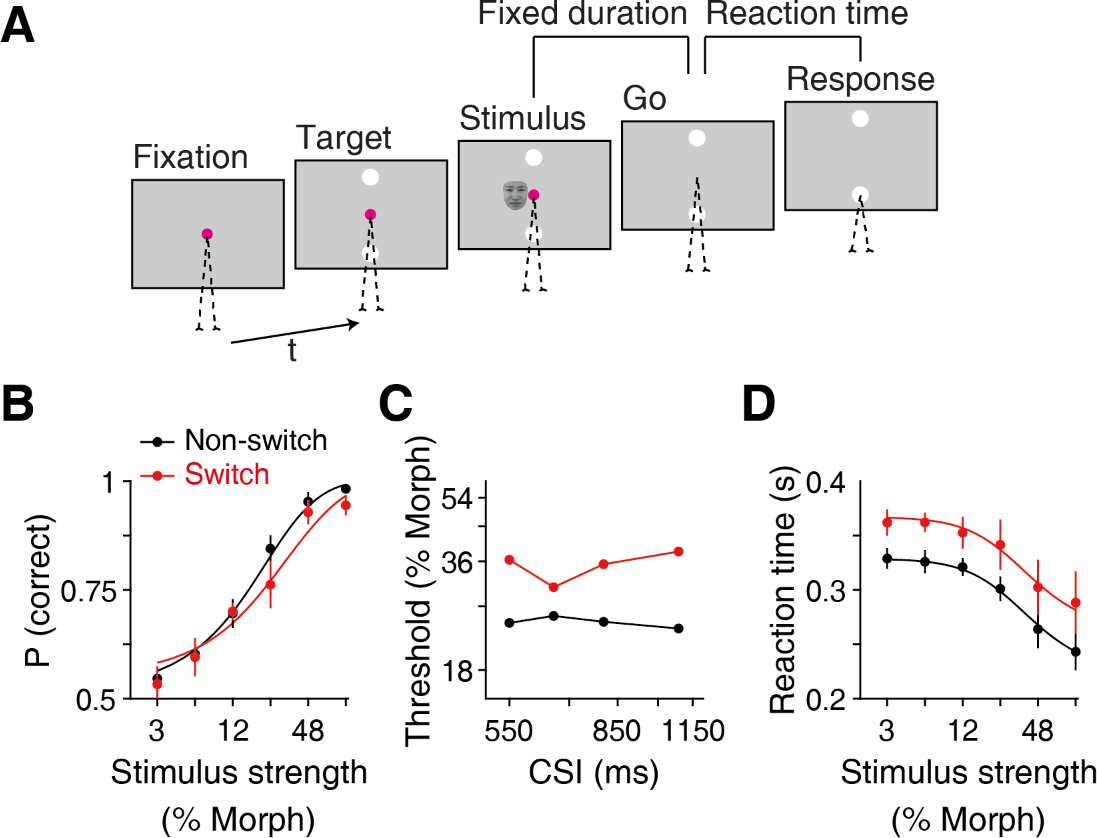
When stimulus duration is limited, task switching affects choice accuracy. (**A**) According to our model (Fig. 3C), participants can maintain choice accuracy after a task switch if they are allowed to accumulate evidence with longer reaction times. If, instead, they are prohibited from accumulating more evidence, the model predicts that task switching directly affects choice accuracy since the reduced weight cannot be compensated. To test this prediction, we performed a version of the face categorization task in which the stimulus duration was fixed (following a truncated exponential distribution; range, 320-640 ms in 106 ms steps). The remaining task parameters were kept identical to those of the main task. Four participants performed this task. (**B**) Reduced choice accuracy was now observed after a task switch, consistent with our prediction. Error bars are S.E.M. across participants. (**C**) The psychophysical threshold (estimated morph level at 81.6% correct rate according to Eq. 6) did not change depending on the duration of the cue-stimulus interval (CSI), consistent with the main task. Trials were divided into four quantiles. (**D**) Reaction times were still longer in the switch trials uniformly across different stimulus strengths. This was also expected from the model.

**Figure 6:**
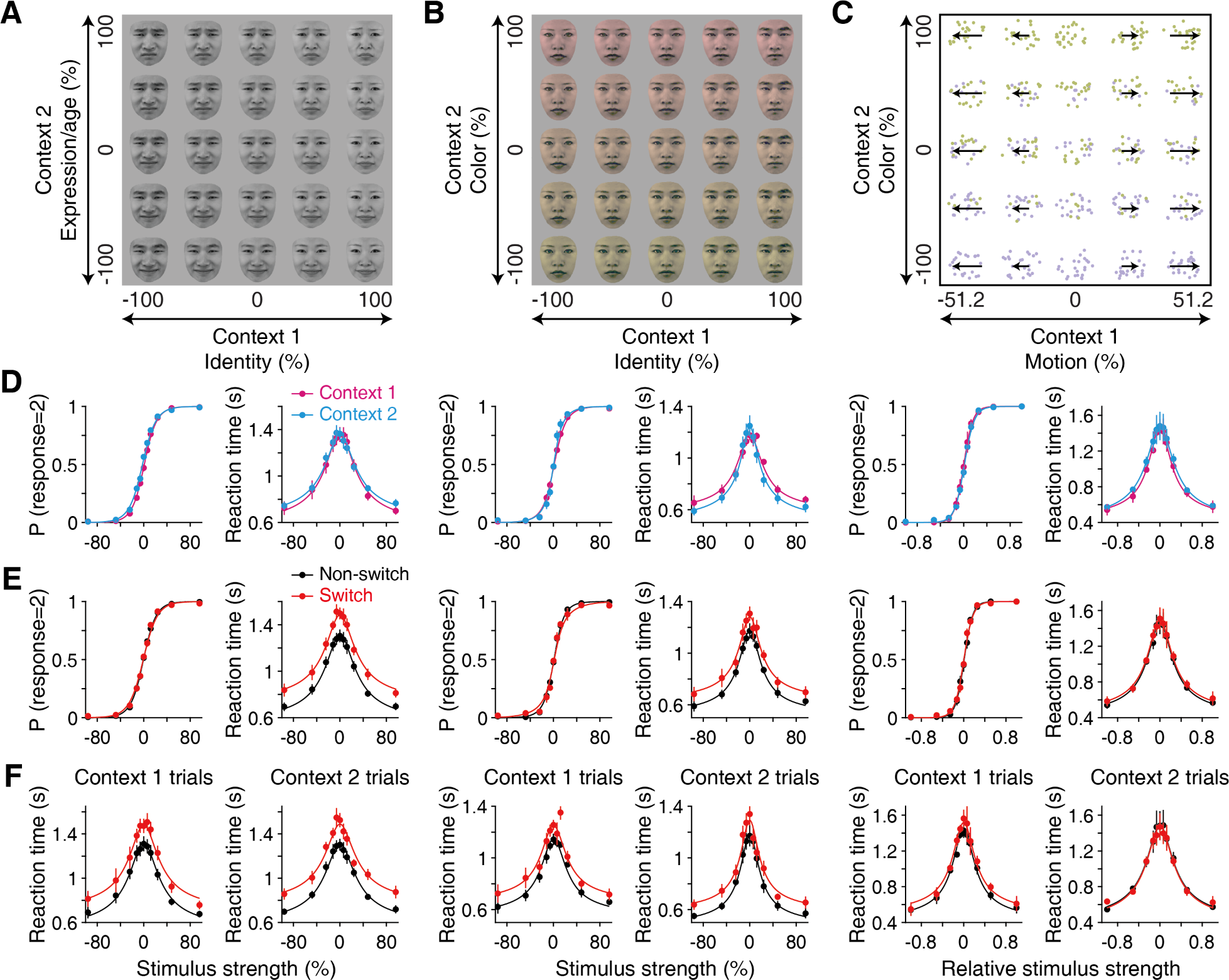
The magnitude of switch cost differs across perceptual tasks. (**A**) The main face task stimuli as in Fig. 1A. (**B**) Facial identity vs. color discrimination task. The identity task was similar to that of the main face task. In the color context, participants categorized the color of face images into red or green. Five participants performed this task. The face images were from the NimStim Face set (Tottenham et al. 2009), but the panel uses images from the Tsinghua Facial Expression Database (Yang et al. 2020) due to copyright issues. (**C**) Color vs. motion direction discrimination task (Mante et al. 2013; Siegel et al. 2015). Participants either judged the net direction of random dot motion (left or right) or judged whether the majority of dots were purple or green in the color context. Three participants performed this task. (**D**) Psychometric and chronometric functions show similar performances across the three tasks and also no apparent difference between the two contexts in each. This ensures that task difficulty or asymmetry of task difficulty between two contexts does not influence switch cost. For the color vs. motion task, the stimulus strengths were normalized by the maximum strength (denoted as “relative stimulus strength”) for proper comparison between the two axes (see Methods section). Error bars indicate S.E.M. across participants. (**E**) The increase in reaction times after rule switch was substantially different among tasks. The conventions are the same as in Fig. 2A. (**F**) The magnitudes of switch cost were similar between the two directions of rule switch in each task. The switch condition in context 1 means the trials immediately after switching from context 2 to 1, while the non-switch condition is all the remaining context 1 trials.

### Switch cost depends on sensory features to be switched

Thus far, the results indicate that task switching reduces the early weighting of sensory information for evidence accumulation, which recovers only after stimulus presentation. A key implication is that the switch cost in our tasks is particularly related to the adjustment in the transformation of sensory information into the decision variable. In this last section, we provide further evidence to support this claim: even under similar experimental structures with similar task difficulty, the magnitude of the switch cost dramatically varies depending on the sensory features that participants were asked to switch.

We compared three different versions of the switch task, including our main reaction-time version of the face categorization task (Figs. 6A-C). We first examined a motion versus color categorization task (Fig. 6C), which has often been used to study context-dependent decision making (Kang et al. 2021; Mante et al. 2013; Siegel et al. 2015). Participants viewed stochastic moving dots colored purple or green and reported either the overall direction of motion or the dominant color. Second, as an intermediate between this and the face categorization task, we examined a face versus color categorization task (Fig. 6B). In this task, participants reported either the identity or the color of the face. In all tasks, stimuli were defined in 2D space, and participants categorized the stimuli based on one of the two axes. The basic task structure, including the frequency of the rule switch and the duration of the CSI, was kept identical across tasks. Furthermore, participants’ overall choice accuracy and RTs were comparable across the tasks (Fig. 6D), and their performance was indistinguishable between the two task rules in each task (accuracy: *p >* 0.088 for all three tasks, two-tailed paired *t*-test; mean RT: *p >* 0.23 for all three tasks). This ensures that the overall task difficulty or imbalance in task difficulty between the two rules (Allport et al. 1994; Yeung and Monsell 2003) does not affect the switch cost.

Despite this carefully tailored comparison, we found substantial differences in the switch costs across the three tasks (Fig. 6E). As demonstrated earlier, in our main face categorization, participants showed an increase of approximately 170 ms in overall RTs across a range of stimulus difficulties. By contrast, the motion versus color task had a much smaller rule switch effect (*∼*31 ms increase in overall RTs; the difference from the main face task: *t*_(9)_ = *−*4.1, *p* = 0.0028, two-tailed *t*-test). The face versus color task had an intermediate level of the rule switch effect (*∼*107 ms increase in overall RTs; the difference from the main face task: *t*_(11)_ = *−*2.1, *p* = 0.056). In all tasks, the effect on choice accuracy was minimal (*α*_1_*_s_* in Eq. 2 was not significant for any of the three tasks; *p >* 0.053, two-tailed *t*-test).

We also found that the increases in RTs were comparable between both directions of the rule switch (i.e., from context 1 to 2 vs. from context 2 to 1) in all three tasks (Fig. 6F; the difference in *β*_0_*_s_* in Eq. 3 between switch directions: *p >* 0.17, two-tailed paired *t*-test). This indicates that the cost was not specific to the switch to particular sensory features, such as facial expressions, but depended more on which pair of features the participants had to switch. For example, the cost of switching from motion discrimination to color discrimination was lower than that of switching from face discrimination to color discrimination (Fig. 6F). Several previous studies have reported altered RTs when switching to the judgment of facial expressions from other judgments, and interpreted the results as the prioritization of biologically significant features (Berger et al. 2019; Elchlepp et al. 2021; Schuch et al. 2012). However, this effect was not observed in our experiments, probably because our face stimuli involved only mild changes in emotion (Fig. 1A).

The dependency of the switch cost on specific perceptual tasks supports the idea that the cost cannot be explained as switching between abstract contextual states in a manner independent of the specific sensory features to be discriminated. Rather, the cost can reflect the difficulty of switching between different sensory readouts, which aligns with our finding of the recovery of sensory processing efficiency after stimulus onset. A comparison of the three tasks alone does not allow us to determine what kinds of sensory features are more difficult to switch between. We speculate that switching features with more overlapping sensory representations can be costly as faces and colors are both encoded in the ventral visual areas (see Discussion section), although it is a formidable challenge to experimentally prove this idea. Nonetheless, the results demonstrate that rule switching is costly not merely because the system requires the transitioning between contextual states.

## Discussion

Humans show a reduction in the accuracy or speed of perceptual decisions after a task rule switch, which has been attributed to top-down cognitive control that requires time to adjust its process (Kiesel et al. 2010; Monsell 2003; Musslick and Cohen 2021). What is puzzling, however, is that even when sufficient time is given, individuals still exhibit substantial switch costs (de Jong 2000; Elchlepp et al. 2021; Li et al. 2019; Meiran et al. 2000; Monsell and Mizon 2006; Rogers and Monsell 1995). We revisited this long-standing observation using recent behavioral measurements and modeling techniques developed to study perceptual decision making (Fetsch 2016; Okazawa et al. 2021b, 2018; Waskom et al. 2019). We found that there was a reduction in the efficiency of processing sensory evidence at the moment of task switching, which recovered within a few hundred milliseconds after stimulus presentation (Fig. 3A). By incorporating this efficiency reduction into an evidence accumulation model, we could accurately explain multiple aspects of the behavioral data in both the switch and non-switch trials (Figs. 3C-G). Furthermore, we found that the cost depended substantially on the type of sensory feature to be switched, even when the task structure remained the same (Fig. 6). We suggest that a critical limitation in perceptual decision making is the flexible switching of the sensory readout, which cannot be fully adjusted based on a context cue alone, but requires the presence of a stimulus to be properly tuned.

Our key finding is attributing the effect of task switching to a specific process that transforms sensory information into decision evidence for accumulation, in contrast to the dominant idea that switch costs reflect the process of switching abstract cognitive states. The cost we observed seems to be unrelated to the processes of recalling a task rule; we found that stimuli with higher strength (e.g., a clearly happy face) along both the task and orthogonal axes neither facilitated nor hindered task switching (Fig. 2A-B), implying that the stimuli themselves (Koch and Allport 2006) or the conjunction of stimuli and context cues (Logan and Bundesen 2004; Schneider and Logan 2005) did not facilitate rule recall. The reduced efficiency appeared to be limited to the first 200-300 ms (Fig. 3A), which corresponds to an early phase of evidence accumulation (Okazawa et al. 2021a; Roitman and Shadlen 2002). Thus, the effect is unlikely to be related to the conflicting action plans between the two rules. Indeed, we observed no congruency effects (Fig. 2K) or interference from task-orthogonal sensory information (Fig. 3B). This is also consistent with recent findings showing a delay in the early components of event-related potentials after a task switch (Elchlepp et al. 2017, 2015). However, this cannot be due to sensory adaptation, priming, or history effects (Kiyonaga et al. 2017; Urai et al. 2019), as we sampled stimuli in an unbiased manner in both tasks. This is also supported by the finding that there was no correlation between the stimulus strength of previous trials and the RT increase (Supplementary Fig. 4). Taken together, we believe that the reduced efficiency is primarily related to the transformation of sensory information into the decision variable.

Then, why does the processing efficiency decrease after task switching for certain perceptual tasks and why does it recover quickly after stimulus onset? While speculative, we hypothesize that this reflects the precision limit of top-down control in adjusting the readout of sensory information to form decision evidence based on context cues alone. After the presentation of a task context cue, decision-making circuits can partially adapt their state to the switched rule so that the task-relevant sensory dimension becomes effective without interference from the previously relevant sensory dimension (see Fig. 3B). However, the circuits cannot optimally tune the readout to convert sensory information into the decision variable with an abstract cue alone; an influx of actual sensory information is needed to guide the circuits to adjust the readout. The difficulty of tuning the readout depends on the specific rules the brain has to switch (Fig. 6). We have yet to specify what factors dictate this difficulty, but we speculate that it is costlier to switch the readout from two overlapping sensory representations such as two face axes or face and color, both of which are encoded in the ventral visual areas (Chang et al. 2017; Chang and Tsao 2017). This interpretation is consistent with previous reports (Elchlepp et al. 2021). For example, switching between visual and auditory tasks shows no residual switch cost (Fintor et al. 2019; Lukas et al. 2010), whereas switching between male and female voice discrimination tasks shows a substantial residual switch cost (Monsell et al. 2019).

This interpretation is different from, but complements, recent compelling theories using RNNs, which propose that switch costs arise because of the extra process of making transitions between two task states in neural spaces (Jaffe et al. 2023; Musslick and Cohen 2021). Such an adjustment of the internal state could delay the onset of decision formation, which might explain the reduced initial sensory efficiency in our task. However, these theories fall short of providing a convincing account of why the recovery of weight must be triggered by stimulus onset rather than by the context cues (Fig. 4F), and why it depends on the specific sensory features to be switched (Fig. 6). We consider rule switching in perceptual tasks to be a multifaceted process that involves both state transition in associative brain areas and adjustment in readout through communication between associative and sensory areas (Okazawa and Kiani 2023).

Our interpretation is also partly related to but distinct from other previously proposed hypotheses regarding the source of the residual switch cost. We highlight the most relevant ideas here. First, some previous studies have suggested that residual switch costs arise because participants occasionally fail to engage and wait until stimulus onset to prepare for a rule switch (de Jong 2000). This claim was made because RTs in switch trials are sometimes as fast as those in non-switch trials, in which participants could be fully prepared. However, such broad distributions of RTs can be produced by noisy evidence accumulation mechanisms without assuming engaged or disengaged states (Fig. 3E). Furthermore, the engagement account does not explain why failure arises in some perceptual tasks but not in others (Fig. 6). Second, several previous studies used the drift-diffusion model to examine switch cost and found effects on non-decision time after a rule switch (Ging-Jehli and Ratcliff 2020; Schmitz and Voss 2012; Schuch and Konrad 2017). Indeed, the patterns of the psychometric and chronometric functions (Fig. 2A) appeared as if there was a change in non-decision time (Fig. 5A). It was only through fine-grained analyses of psychophysical reverse correlations that we correctly attributed them to the initial reduction in sensory weighting (Fig. 3). Finally, it has been proposed that humans have limited ability to shift their attention to a relevant stimulus dimension before a stimulus appears (Elchlepp et al. 2017, 2015, 2021). This idea is most relevant to our hypothesis, but we found that stimuli with higher strengths, which likely attracted stronger attention to the relevant dimension, did not facilitate task switching (Fig. 2A). Therefore, we interpret this as the cost of adjusting the sensory information flow instead of attention.

This study identified a specific process that gives rise to switch costs, but it should be noted that switch costs likely result from multiple factors whose relative contributions depend on task details (Kiesel et al. 2010; Koch et al. 2010, 2018; Monsell 2003; Monsell and Mizon 2006; Ruge et al. 2013). For example, studies using shorter CSIs than ours have identified part switch cost that was strongly dependent on the CSI (Meiran 1996). This likely reflects a process more directly related to internal preparation based on contextual cues (Jaffe et al. 2023; Musslick and Cohen 2021). On a related note, when there are multiple cues for one context, a cue change alone without task switching degrades behavioral performance, known as the cue switch cost (Arrington and Logan 2004; Logan and Bundesen 2004; Mayr and Kliegl 2003; Schneider and Logan 2005). Such cue-encoding mechanisms are beyond the scope of our decision-making models. Regarding response encoding, the congruency effect or response conflict is often observed along with switch costs (Hyafil et al. 2009; Monsell et al. 2000; Yeung and Monsell 2003), whereas our behavioral results lacked these effects (Fig. 2K). These are typically associated with tasks using firmly established associations between stimuli and responses, such as the Stroop task (Allport et al. 1994). The lack of these effects in our design might stem from arbitrary associations between face stimuli and saccade directions. We expect that the diverse effects of task switching observed in previous studies can be investigated by extending our quantitative modeling framework to different task structures.

This study focused on switching between perceptual tasks, but similar principles may apply to other cognitive tasks. The residual switch cost has been reported in many non-perceptual tasks, such as number or lexical categorizations (Monsell 2003). Although the sensory readout may not be a major bottleneck in these tasks, the process of converting sensory inputs into decision evidence may require fine adjustments in circuit computations. Although top-down control can partially align these circuit computations with a given task demand, detailed computations may require further adjustments after a stimulus is presented and the circuits start to operate, leading to limited behavioral performance unique to biological neural networks.

## Methods

### Participants and experimental setup

Sixteen human participants (20-40 years old, 6 males and 10 females, students or employees at the Chinese Academy of Sciences) were recruited in total for the experiments. We collected a large number of trials from each participant (total 54,938 trials; *∼*1,500-3,000 trials per participant for each experiment) to perform psychophysical reverse correlation and model fitting (Ahumada Jr 1996; Smith and Little 2018). Each participant additionally underwent extensive training (*∼*2,000 trials) before the main data collection. The participant count in each experiment was comparable to those in previous studies using similar approaches (Kang et al. 2021; Levi et al. 2018; Stine et al. 2020). All the participants had normal or corrected-to-normal vision and were näıve to the purpose of the experiment. Written informed consent was obtained from all participants. All experimental procedures were approved by the Institutional Review Board of the Center for Excellence in Brain Science and Intelligence Technology, Institute of Neuroscience, Chinese Academy of Sciences.

Throughout the experiments, the participants were seated in a height-adjustable chair in a semi-dark room with their chin and forehead supported by a tower-mounted chinrest. The chinrest had a fixed position to ensure a stable viewing distance (57 cm) from the CRT display monitor (17-inch IBM P77 and 21-inch SUN GDM-5010P; 75 Hz refresh rate; 1024 *×* 768 pixels screen resolution).

Stimulus presentation was controlled using the Psychophysics Toolbox (Brainard and Vision 1997) and MATLAB (MathWorks, MA, USA). Eye movements were monitored using a high-speed infrared camera (Eyelink; SR Research, Ottawa, Canada). The gaze position was recorded at 1 kHz.

### Task designs

#### Context-dependent face categorization task

To investigate flexible task switching, we designed a context-dependent face categorization task. We chose face categorization because previous studies successfully explained behavior using a simple evidence accumulation model with psychophysical reverse correlation (Okazawa et al. 2021b). Furthermore, face stimuli can be naturally categorized along multiple sensory dimensions such as identity, expression, and age, making them suitable for studying flexible rule switching.

Participants categorized faces defined in a 2D face space (Fig. 1A) according to one of two categorization rules. Categorization rules were switched every 2-6 trials within the experimental blocks and were indicated by the color of a fixation point such that participants were always informed of the rule. The two rules were facial identity versus expression categorization for six of the eight participants who participated in this experiment, and facial identity versus age categorization for the remaining two of the eight participants. We used these two conditions to ensure that the switch cost effects were not due to specific types of facial features such as expressions (Elchlepp et al. 2021). For the identity rule, the participants categorized faces into one of two facial identities. For the expression and age rules, the participants categorized faces as happy/sad or old/young. As the behaviors of these two groups were comparable (Supplementary Fig. 1), we averaged the results in the main section.

Each trial began when participants fixated on a fixation point at the center of the screen (0.5° diameter). The color of the fixation point was either cyan or magenta and cued one of the two categorization rules. After a short delay (150-300 ms, truncated exponential distribution), two white target dots appeared 7° above and below the fixation point. Shortly thereafter (300-1200 ms, truncated exponential distribution), a face stimulus (size, *∼* 4° *×* 4°) appeared on the screen parafoveally (stimulus center, 1.5° to the left of the fixation point). We placed the stimuli parafoveally to encourage participants to judge the face stimulus as a whole, rather than focusing on local features. Participants reported the category of the presented face by making a saccade to one of the two targets whenever they were ready (reaction-time task). Associations between face categories and target positions were counterbalanced across participants in each context. The stimulus was extinguished immediately after saccade initiation. If the participants did not make a decision within 5 s, the trial was aborted (*<* 0.2% of the trials). Distinct auditory feedback was delivered for correct and incorrect choices. When the face was ambiguous (halfway between the two prototypes on the morph continuum), the correct feedback was delivered in a random half of the trial. Following feedback, the next trial began after a 1.2 s inter-trial interval.

We created a 2D face space by continuously morphing four prototype faces. The prototype faces were obtained from the Tsinghua Facial Expression Database (Yang et al. 2020) and the NimStim Face set (Tottenham et al. 2009), which contains photographs of the same identities with different expressions. To create young/old prototypes, we used free software (Alaluf et al. 2021) that synthesizes younger or older faces from an original photograph. Morphed facial images were created from the prototype faces using a custom program (Okazawa et al. 2021b). The program linearly interpolates the positions of manually defined anchor points on the facial images and the textures inside the tessellated triangles defined by the anchor points. This algorithm can also independently morph different facial features (eyes, nose, and mouth).

Two stimulus axes were generated from four prototypes (e.g., images of happy person A, sad person A, happy person B, and sad person B), but we took extra caution in making the two axes orthogonal (factorial) (Folstein et al. 2012). For example, the morph axis connecting the happy and sad faces of identity A (*A_H_, A_S_*) is not equivalent to the morph axis connecting the happy and sad faces of identity B (*B_H_, B_S_*). The factorization of the two axes requires the construction of the following two morph vectors and morphing faces along these two axes:

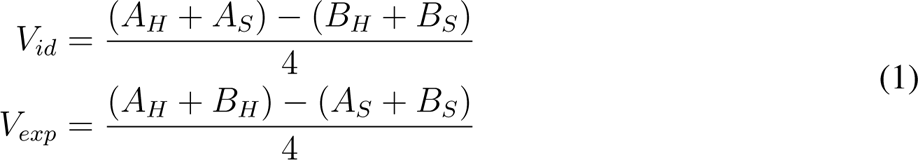

The center of the face space (0% morph level) is the average of all four faces (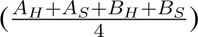) and *±*100% morph levels for each stimulus axis correspond to the addition or subtraction of the above vectors to the average face. On each trial, we sampled one morph level from the following 11 levels for both axes regardless of the categorization rules: −96%, −48%, −24%, −12%, −6%, 0%, +6%, +12%, +24%, +48%, +96%. As shown in Supplementary Figures 1A-B, the participants had roughly equal discriminability along the two axes.

We added random temporal fluctuations in morph levels to the sampled value in each trial to examine how participants weighted the evidence conferred by the face stimuli over time (i.e., psychophysical reverse correlation). The morph level was randomly sampled every 106.7 ms (eight monitor frames) from a Gaussian distribution with an SD of 20%. This fluctuation duration provided us with sufficiently precise measurements of participants’ weighting characteristics in their *∼*1 s decision time, while the duration was long enough to ensure a subliminal transition of morph levels. Between the two morphed face images, we interleaved a noise mask (phase randomization of 0% morph face) with a smooth cosine transition function during the eight monitor frames (Okazawa et al. 2021b). This mask prevented participants from noticing fluctuations in morph levels during stimulus presentation. Random fluctuations were applied independently to each facial feature (eyes, nose, and mouth) along both the task and orthogonal axes while keeping the average morph level constant within a trial. Although the independent fluctuations across facial features allowed us to examine the spatial weighting of evidence (Okazawa et al. 2021b), most of our analyses used the average morph levels of the three features because our primary goal was to test temporal weighting. The psychophysical kernels for the individual features are shown in Supplementary Figure 5.

We collected data from eight participants for this task (24,358 trials in total; 3,045 *±* 353 trials per participant). Prior to the main data collection, the participants underwent extensive training (on average, 5 sessions and 2,200 trials) to ensure stable behavioral accuracy under both rules.

#### Fixed stimulus duration task

In the RT task described above, participants showed longer RTs in the switch trials, but their choice accuracy was maintained, possibly because they were able to collect more evidence with additional RTs. Therefore, we designed an alternative task in which we limited the stimulus duration to test whether choice accuracy deteriorated when the collection of additional evidence was prohibited.

In each trial, a stimulus was presented for a variable duration (truncated exponential distribution; range, 320-640 ms, in steps of 106.7 ms; mean, 416 ms). This distribution has a flat hazard rate and minimizes the participants’ ability to anticipate the end of a stimulus (Ghose and Maunsell 2002). The fixation point disappeared with stimulus termination as the go cue. Participants had to select a target within 0.7 s after the go cue (the proportion of timeout trials: *<* 7% for all participants). The RT in this task was defined as the time interval between the stimulus offset and saccadic response. The remaining task parameters, including the stimulus design and duration of the CSI, were identical to those in the RT task.

Four participants performed this fixed duration task (12,703 trials in total; 3176 *±* 356 trials per participant). Three participated in the RT task with the same facial stimulus set.

#### Motion versus color and face versus color categorization tasks

To test whether similar degrees of switch costs occur with perceptual tasks using simpler sensory features, we performed motion versus color categorization of colored stochastic moving dots (Kang et al. 2021; Mante et al. 2013; Siegel et al. 2015; Fig. 6C). In the motion task, participants reported whether the dots were overall moving to the left or right by making a saccade to one of the two targets positioned to the left or right of the fixation point. In the color task, participants reported whether the majority of the dots were purple or green by choosing one of the same two targets. The two targets were colored purple and green to indicate the association. The overall trial structure was similar to that of the RT version of the face task. To avoid interference with color judgment, task contexts were conveyed by the shape of the fixation point rather than by its color (white triangle or cross). A random dot stimulus appeared within a 6-degree circular aperture centered at the fixation point. It consisted of three independent sets of moving dots displayed in consecutive frames (Britten et al. 1992). Each set of dots was shown for one video frame and then replotted three frames later (Δ*t* = 40 ms; density, 16.7 *dots/deg*^2^*/s*). When replotted, a subset of dots was offset from their original location (speed, 5 °*/s*), whereas the remaining dots were placed randomly. The colors of the dots were chosen to be equiluminant green (L = 20.0, x = 0.386, y = 0.494) or purple (L = 20.0, x = 0.224, y = 0.182).

The stimulus strength of the motion (motion coherence) was defined as the percentage of dots moving coherently in the correct direction. The stimulus strength of the color (color coherence) was defined as the difference between the percentage of green and purple dots (Kang et al. 2021; Mante et al. 2013). For example, +100% color coherence meant all the dots were green, −100% color coherence meant all the dots were purple, and 0% color coherence meant that green and purple dots were equally likely to be present. On each trial, a motion coherence was chosen from the following set: −51.2%, −25.6%, −12.8%, −6.4%, −3.2%, 0%, +3.2%, +6.4%, +12.8%, +25.6%, +51.2%. A color coherence was chosen from a different set to match the difficulty with the dots task: −100.0%, −51.2%, −25.6%, −12.8%, −6.4%, 0%, +6.4%, +12.8%, +25.6%, +51.2%, +100.0%.

For visualization purposes (Figs. 6D-F, right panel), we scaled the eleven stimulus strengths of each task to range from *−*1 to 1 and denoted it as “relative stimulus strength”. We also performed a face versus color categorization task as a control (Fig. 6B). In this task, participants categorized a colored facial stimulus according to its facial identity or color. As in the motion versus color task, the task contexts were indicated by the shape of the fixation point (white triangle or cross). Two prototype facial identities were chosen from the NimStim Face set (Tottenham et al. 2009), and the faces were uniformly colored with a value in the CIE-1931 xy color space linearly interpolated between the two prototype colors. Because we found that the participants had different color-discrimination thresholds, we chose different prototype colors for different participants to match the task difficulty between the face and color contexts. These prototypes were red (set 1: x = 0.374, y = 0.274; set 2: x = 0.371, y = 0.318; set 3: x = 0.369, y = 0.339) and green (set 1: x = 0.361, y = 0.443; set 2: x = 0.365, y = 0.400; set 3: x = 0.366, y = 0.378). The luminance of each image pixel was kept constant. The stimulus strength of color was defined as the distance from the prototypes; each prototype corresponded to −100% and 100% strength, and the intermediate values were their linear interpolation. We had 11 levels: −96%, − 48%, −24%, −12%, −6%, 0%, +6%, +12%, +24%, +48%, +96%. To approximate the fluctuations in sensory evidence that occurred in the other tasks, we introduced a random variation in color to a stimulus. The color strengths were randomly sampled from a Gaussian distribution with an SD of 20% and updated every 13.3 ms (one monitor frame). This rapid fluctuation mimicked the stochasticity of color strength in the motion versus color task, where the color of each dot was resampled in every monitor frame.

Three participants performed the motion versus color task (8,755 trials in total; 2,918 *±* 56 trials per participant), and five participants performed the face versus color task (9,122 trials in total; 1,824 *±* 252 trials per participant). One participant also performed the main face task. Participants received extensive training for each task before main data collection (on average, 4 sessions and 2,000 trials).

### Data analysis

#### Psychometric and chronometric functions

Throughout the analyses, we defined the trials immediately after a rule switch as switch trials and the rest of the trials as non-switch. The first trial of each experimental block was excluded. We confirmed that history effects such as post-error slowing (Purcell and Kiani 2016) did not affect our conclusions (Supplementary Fig. 4).

To quantify the differences in the participants’ behavioral performance between the switch and non-switch trials, we fitted the following logistic function to the choice data for each participant:

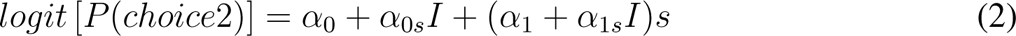

where *logit*(*p*) = *log*(*p/*(1 *− p*)), *s* is the morph level (ranging from –1 to 1) and *I* is an indicator variable that is 0 for non-switch trials and 1 for switch trials. *α*_0_*, α*_0_*_s_, α*_1_ and *α*_1_*_s_* are regression coefficients reflecting choice bias in non-switch trials, the difference in choice bias between non-switch and switch trials, accuracy in non-switch trials, and the difference in accuracy between non-switch and switch trials, respectively.

The difference in the mean RTs between the switch and non-switch trials was evaluated using a hyperbolic tangent function:

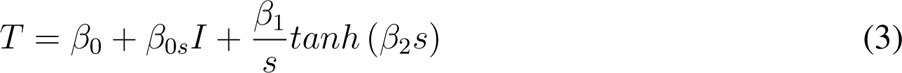

where *T* denotes mean RTs. *β*_1_*, β*_2_ are regression coefficients that reflect the dependency of RTs on stimulus strength for non-switch trials, and *β*_0_ is a stimulus-independent term. Because the increase in RTs for switch trials was independent of stimulus strength (Fig. 2C), we added *β*_0_*_s_* to account for the RTs in switch trials.

#### Psychophysical reverse correlation

To quantify the effect of stimulus fluctuations on choice, we performed a psychophysical reverse correlation (Ahumada Jr 1996; Okazawa et al. 2018). Psychophysical kernels *K*(*t*) were calculated as the difference in the average fluctuations of the morph levels, conditional on the participants’ choices, as follows:

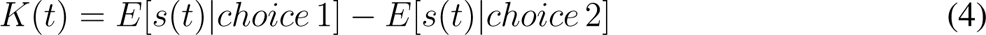

where *s*(*t*) represents the morph level of the facial stimulus at time *t*. Although the morph levels fluctuated independently for the three facial features (eyes, nose, and mouth), we averaged them for each stimulus frame to focus on the effects of temporal fluctuations. When averaging, we weighted the fluctuations of each feature according to the fitted sensitivity parameters of that feature in the drift-diffusion model (*k_e_, k_n_, k_m_* in Eq. 7; see below) so that a more informative feature makes a larger contribution to the kernel. The psychophysical kernels for the individual features are shown in Supplementary Figure 5, and are consistent with our main conclusions. We used trials with low average morph levels (average level 0-12%), in which there was a sufficient number of trials for both choices. For trials with nonzero average morph levels, the average was subtracted from the fluctuations. We used fluctuations up to the median RT aligned to the stimulus or saccade onset to ensure that at least half of the trials contributed to the kernels at all times. Three-point boxcar smoothing was applied to the kernels for denoising.

#### Testing the effect of cue-stimulus interval on behavior

Our tasks had a variable delay between the onset of participants’ fixation—when they recognized the current task rule based on the fixation color—and stimulus presentation (range, 0.5-1.5 s; median, 0.72 s; roughly followed truncated exponential distribution but depended on participants’ fixation onset; Fig. 2H). This cue-stimulus interval (CSI) allowed participants to prepare for rule switching. To test the effect of CSI on RTs, we divided trials into seven quantiles based on CSIs and calculated the difference in RTs between the switch and non-switch trials (ΔRT) for each quantile. ΔRT was quantified as *β*_0_*_s_* in Eq. 3, because it was largely independent of the stimulus strength (Fig. 2C). A direct comparison of RTs between switch and non-switch trials yielded similar results. We performed a linear regression between CSI and *β*_0_*_s_* for each participant, then averaged the results across participants (Fig. 2J).

The effect of the CSI on the reduction in psychophysical kernels was examined by dividing the trials into two groups based on the median CSI. We focused on the kernel of the first stimulus frame and defined the reduction in kernel amplitude for the switch trials as

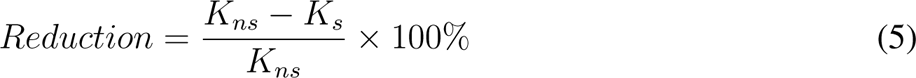

where *K_s_*and *K_ns_* are the kernels of the switch and non-switch trials, respectively, calculated as described above for each participant.

#### Discrimination threshold in the fixed stimulus duration task

To test whether the choice accuracy was affected by CSIs in the fixed stimulus duration task, we plotted the discrimination thresholds of the switch and non-switch trials, each divided into four quantiles of CSIs (Fig. 5C). For each quantile, we fit the following logistic function to the choice data,

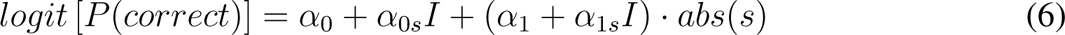

From the fitted curve, we calculated the discrimination threshold, which is defined as the stimulus strength at 81.6% correct (Kiani et al. 2008).

### Model fit and evaluation

To examine the computational mechanisms underlying the behavioral differences between the switch and non-switch trials, we fitted several variants of evidence accumulation models to the data. Previous studies showed that face categorization behavior without a rule switch can be accurately explained by a drift-diffusion model that linearly accumulates spatiotemporal evidence (Okazawa et al. 2021b, 2018). Therefore, we fitted this model to non-switch trials and attempted to explain the switch trials using additional mechanisms.

#### Drift-diffusion model for non-switch trials

To fit the non-switch trials, we employed a model previously developed for face categorization behaviors (Okazawa et al. 2021b, 2018). The model receives spatiotemporal fluctuations in morph levels, linearly accumulates evidence toward an upper or lower bound, and commits to the choice associated with the bound when it is reached. RT is the sum of the time required to reach a bound and the additional non-decision time (Fig. 3C).

In our task, the morph levels of the three facial features (eyes, nose, and mouth) fluctuated along both the task and orthogonal axes, resulting in six morph levels. However, because participants rarely confused the task rule (Fig. 2A) and did not show influences from task-orthogonal information (Fig. 2B), we assumed that only the task axes contributed to forming the momentary evidence (*µ*(*t*)) in each context:

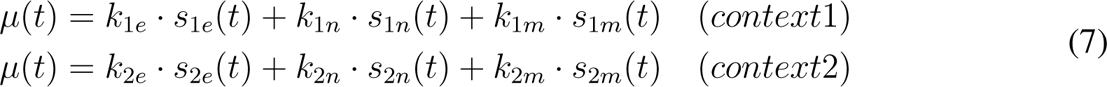

where *s*_1_*_e_, s*_1_*_n_, s*_1_*_m_, s*_2_*_e_, s*_2_*_n_, s*_2_*_m_* are the morph levels of the individual facial features along the task axis, and *k*_1_*_e_, k*_1_*_n_, k*_1_*_m_, k*_2_*_e_, k*_2_*_n_, k*_2_*_m_* are the sensitivities associated with them. The sensitivities for each feature were parameterized independently for each context because facial features can be weighted differently for different tasks (Okazawa et al. 2021b; Schyns et al. 2002).

Momentary evidence is then accumulated over time to form the decision variable (*v*):

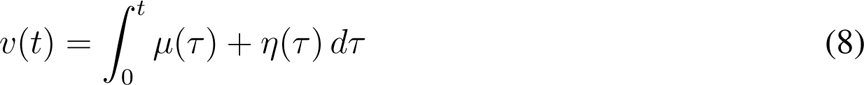

where *η*(*τ*) represents internal (neural) noise in the sensory, inference, or integration processes, assumed to follow a Gaussian distribution with mean 0 and SD *σ*(*t*). Under these assumptions, the probability that the decision variable has value *v* at time *t* satisfies the Fokker-Planck equation

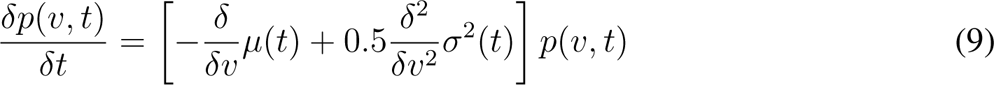

where *p*(*v, t*) denotes the probability density. The accumulation process started from zero evidence and continued until the decision variable reached one of the two boundaries (*±B*) indicating two choices. Thus, the above partial differential equation has initial and boundary conditions as follows:

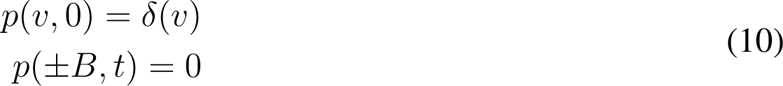

where *δ*(*v*) denotes the Dirac delta function. The diffusion noise (*σ*(*t*)) was set to 1 and the bound and drift rate were defined in a unit of diffusion noise. The bound-crossing time is convolved with the distribution of non-decision time, which was defined as a Gaussian distribution with a mean of *T*_0_ and an SD of *σ_T_*_0_, to calculate the RT. The SD, *σ_T_*_0_, was always set to one-third of *T*_0_ to reduce the number of the free parameters (Churchland et al. 2008).

In total, the model had eight free parameters (*k*_1_*_e_, k*_1_*_n_, k*_1_*_m_, k*_2_*_e_, k*_2_*_n_, k*_2_*_m_, B, T*_0_) to fit the non-switch trials. The majority of the parameters accounted for the sensitivities of the three facial features in each rule, and by averaging the fluctuations of the three features, the model could be reduced to four parameters (*k*_1_*, k*_2_*, B, T*_0_; one sensitivity parameter for each task context). This reduced model performed equally well in fitting the behavioral data and yielded similar results, except that it did not account for feature sensitivities. Nevertheless, we used the 8-parameter model for the main results to conform to our previous study (Okazawa et al. 2021b). The model had the same bound height (*B*) and non-decision time (*T*_0_) in the two contexts. This was justified by the fact that the participants had similar psychometric and chronometric functions in both contexts (Supplementary Figs. 1A-B).

We fitted the model to the participants’ choices and RT distributions using maximum likelihood estimation. Given a set of parameters, the stimulus fluctuations in each trial were used to calculate the RT distributions of the two choices according to the aforementioned model formulation. From these distributions, we derived the likelihood of observing the participants’ choices and RTs. Summing the likelihoods across all the trials yielded the total likelihood of the parameter set. We used a simplex search method (*fminsearch* in MATLAB) to determine the parameter set that maximized the summed likelihood. To avoid local maxima, we repeated the fitting process using multiple initial parameter sets and selected the largest overall likelihood as the final result. Fitting was performed for each participant, and included trials with all stimulus strengths. The fitting curves shown in Figures 3 and 4 represent the averages across the participants.

#### The reduction in initial sensory weighting in switch trials

To account for the reduced effect of initial stimulus fluctuations in the switch trials (Fig. 3A), we added a dynamic sensory weighting function to the drift-diffusion model (Levi et al. 2018; Okazawa et al. 2018). Assuming that sensory weight drops after a task switch and recovers gradually during stimulus presentation, we modeled the sensory weight at each time *t* (*w*(*t*)) in the switch trials as a ramp function:

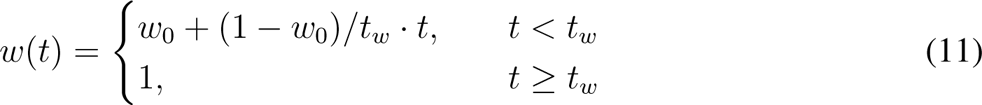

where *w*_0_ is the initial value and *t_w_* is the time required for the weight to return to baseline. This weighting was constant in non-switch trials.

Sensory weighting can affect evidence accumulation in two ways. First, it can modulate only the signal component (i.e., drift rate) of the accumulation. This can be formulated by extending Eq. 8 as:

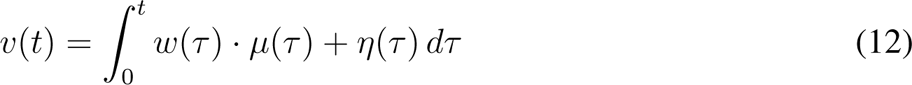

Second, it can modulate the diffusion noise along with the drift rate:

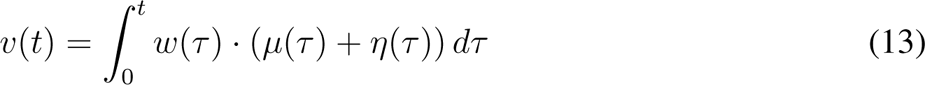

These two forms make different assumptions regarding the noise source. If noise arises during the accumulation process, the weighting function does not affect the noise (Eq. 12), but if noise arises in the sensory process or in the process of converting sensory information into evidence, weighting can also be applied to the noise (Eq. 13). In Supplementary Figure 2, we tested both forms and found that the latter explained the behavioral data well. This is consistent with previous findings that noise in perceptual decision making is largely due to sensory or inference processes (Brunton et al. 2013; Drugowitsch et al. 2016; Waskom and Kiani 2018). Therefore, we use the latter form in our main results.

To fit this model to the switch trials, we changed only the parameters for weight reduction (*w*_0_*, t_w_*) during the maximum likelihood estimation for individual participants. All other parameters were fixed at the values fitted to the non-switch trials. This poses strong constraints on the model because only two parameters were used to account for behavioral changes in the switch trials. We also confirmed that the model parameters converged to similar values when all parameters were simultaneously fitted to the switch trial data.

#### Alternative models

To examine whether other mechanisms could explain the observed switch costs, we simulated a variety of alternative models. Because sensitivity (*k*), bound height (*B*), and non-decision time (*T*_0_) in the drift-diffusion model affect RTs, we first examined whether any of these parameters accounted for the data in the switch trials. Similar to the model with initial weight reduction, we started with the model parameters fitted to the non-switch trials, and then allowed the model to adjust these parameters to fit the switch trials using maximum likelihood estimation (non-decision time: Figs. 4A-C; sensitivity and bound height: Supplementary Figs. 3C-E).

We also tested whether a leak in evidence accumulation explains the results (Supplementary Fig. 3B), because it increases RTs and leads to lower kernel amplitudes early in trials owing to the gradual loss of information over time. The drift diffusion model with a leak rate (*λ*) becomes an Ornstein-Uhlenbeck process (Bogacz et al. 2006), whose Fokker-Planck equation is

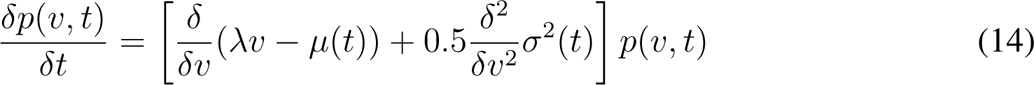

A larger leak rate indicates a greater loss of information over time.

To further examine whether task preparation after receiving the context cue interfered with evidence accumulation and led to the observed switch cost, we extended our model to include task preparation time (Fig. 4D). This extended model assumes that, after participants fixated on the fixation point indicating a rule switch, they needed a fixed amount of time (modeled as a Gaussian distribution with mean *P_m_* and SD *P_sd_*) to prepare for the switch. Since evidence accumulation starts only after this preparation time, an early period of the stimulus sequence does not influence the participants’ decisions if this period overlaps with the preparation time. The key difference from the main model is that the duration of this ineffective stimulus period depends on the CSI, which varied across trials in our task (Fig. 2H). The model was allowed to adjust *P_m_* to fit the switch trials using the same maximum likelihood estimation. The SD, *P_sd_*, was maintained at one-third of *P_m_*, because we found that *P_sd_*takes an extremely large value when fitted as a free parameter to account for the switch cost in trials with long CSIs (Fig. 4F), making the model inappropriate as a hypothesis for task preparation.

#### Generation of model psychophysical kernels and RT distributions

Because the above model formulation specifies choices and RTs, but not psychophysical kernels, we relied on Monte Carlo simulations to estimate the model kernels. We created 10^5^ simulated trials with 0-12% morph levels with the same morph fluctuation parameters as in the main task (SD, 20%), and generated the responses of the fitted models to these stimulus patterns. We then used the model choices and RTs to calculate their psychophysical kernels, as we did for the human data (thick gray lines in Figs. 3F-G, 4C, and 4E). Thus, the model kernels were not directly fitted to the participants’ kernels, but were generated from an independent set of stimulus fluctuations, making the comparison of data and models informative. Similarly, the RT distributions of the models (Fig. 3E) were generated from simulations with an independent set of morph fluctuations to ensure an accurate comparison of the data and models.

## Acknowledgements

We thank Roozbeh Kiani, Bin Min, Yiteng Zhang, and Tianming Yang for their helpful discussions and comments on earlier versions of the manuscript. We thank Jiahao Wu, Chang Liu, and Mingrui Zheng for their assistance with data collection. This work was supported by the National Science and Technology Innovation 2030 Major Program (grant 2021ZD0203703) and the National Natural Science Fund for Excellent Young Scientists Fund Program (overseas).

## Data and code availability

Data and code used in this study will be made publicly available on GitHub upon official publication.

---

**Figure S1:**
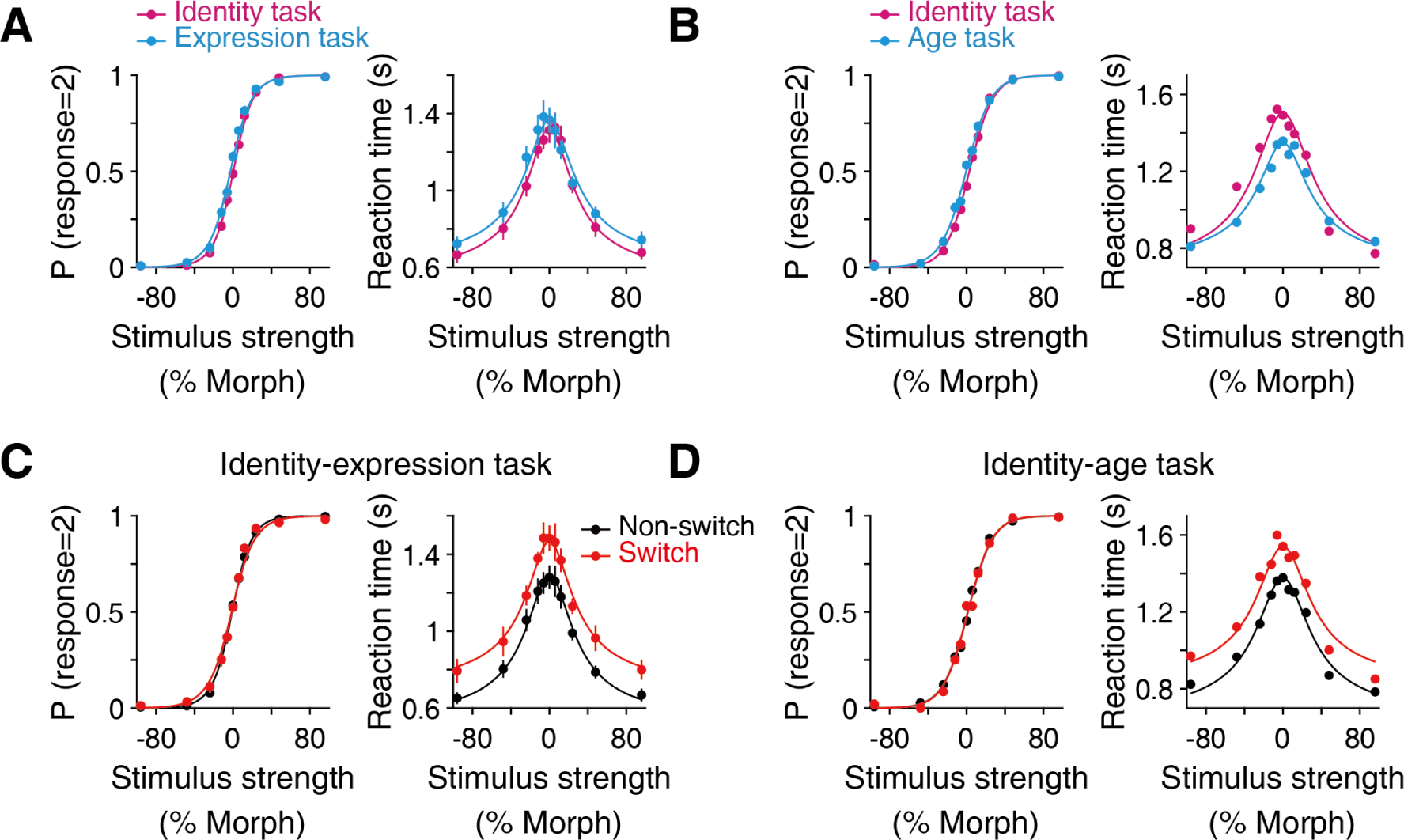
Choices and reaction times for different versions of flexible face categorization task. (**A, B**) In the main face categorization task (Fig. 1), six participants performed identity vs. expression categorization (A), and two performed identity vs. age categorization (B). The patterns of choices and reaction times were similar between these conditions and also nearly indistinguishable between the two categorization rules in each condition. (**C, D**) In both conditions, we observed an increase in reaction times with little change in choice accuracy after rule switch. Since the patterns and magnitudes of switch cost were comparable between the conditions, we averaged the two tasks in the main figures (Figs. 2-4).

**Figure S2:**
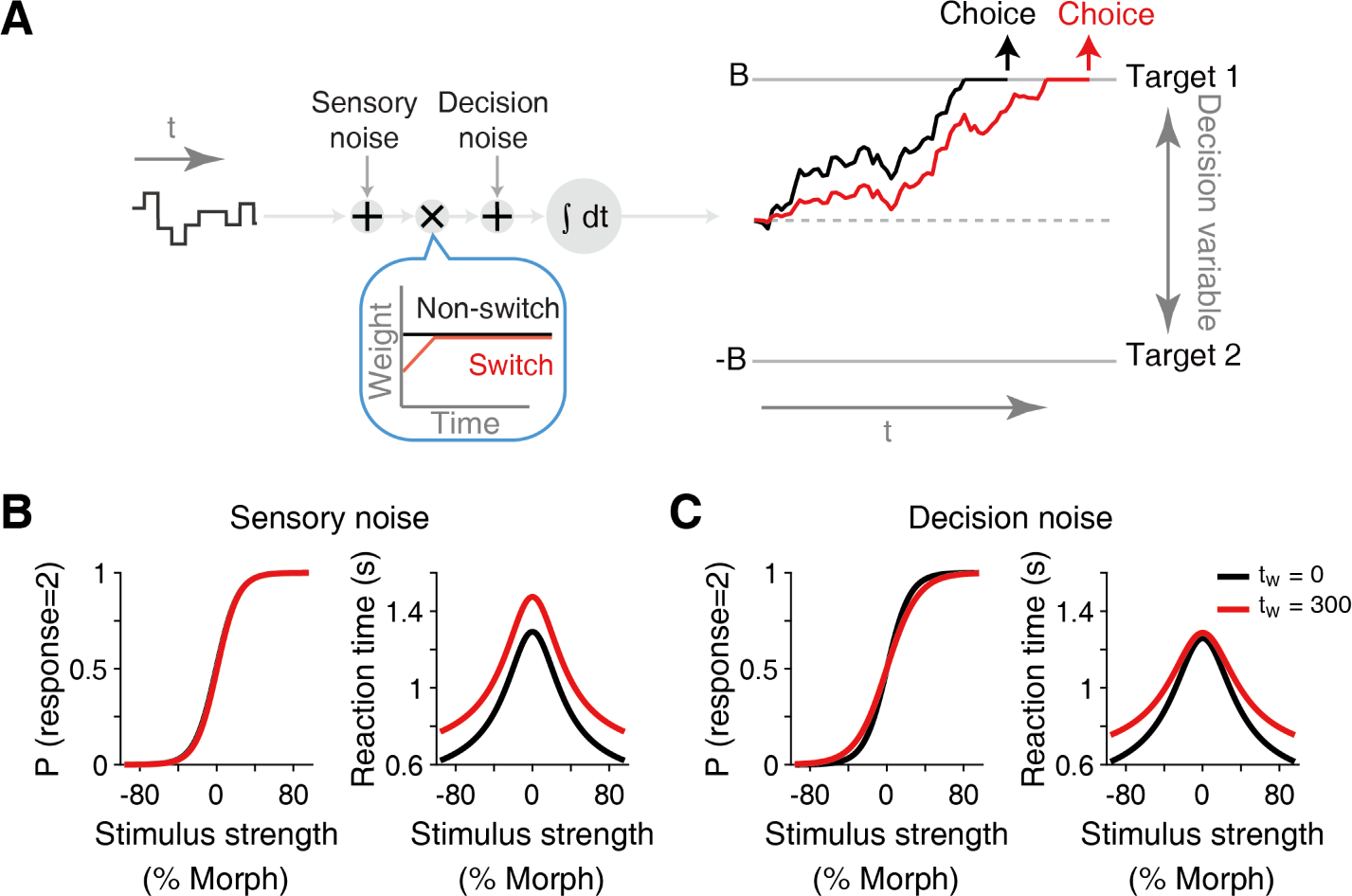
Two different forms of the model with dynamic sensory weighting functions. (**A**) In the evidence accumulation model, the momentary sensory evidence is accumulated along with internal (neural) noise (η in Eq. 8). The sensory weighting function can modulate the evidence either before or after this noise is added. These two expressions (Eq. 12 and Eq. 13) have different assumptions regarding the source of the noise. If the source of the noise is considered to be in the sensory processes or in the process of converting sensory input into decision evidence (inference process), it is natural to assume that the sensory weighting also applies to the noise. If the noise arises in the process of evidence accumulation, the sensory weighting will not apply to it. (**B, C**) These two model expressions yield different outcomes. The plots are based on Monte Carlo simulations performed with an arbitrary set of model parameters for demonstration purposes (k = 0.15, B = 30, mean of T_0_ = 400, SD of T_0_ = 133, w_0_ = 0, t_w_ = 0 or 300 for black and red lines respectively). When the weighting is applied to the noise, both the signal and noise are reduced early in the evidence accumulation. Thus, the model shows longer reaction times for all stimulus strengths. By contrast, when the weighting does not apply to the noise, only the signal is reduced, and thus the model shows longer reaction times only for stronger stimulus strengths. Our empirical data were largely consistent with the former pattern. Therefore, for simplicity, we used the former model in our main results. This model is also consistent with previous reports that noise in perceptual decision making arises predominantly in the sensory or inference processes (Brunton et al. 2013; Drugowitsch et al. 2016; Waskom and Kiani 2018).

**Figure S3:**
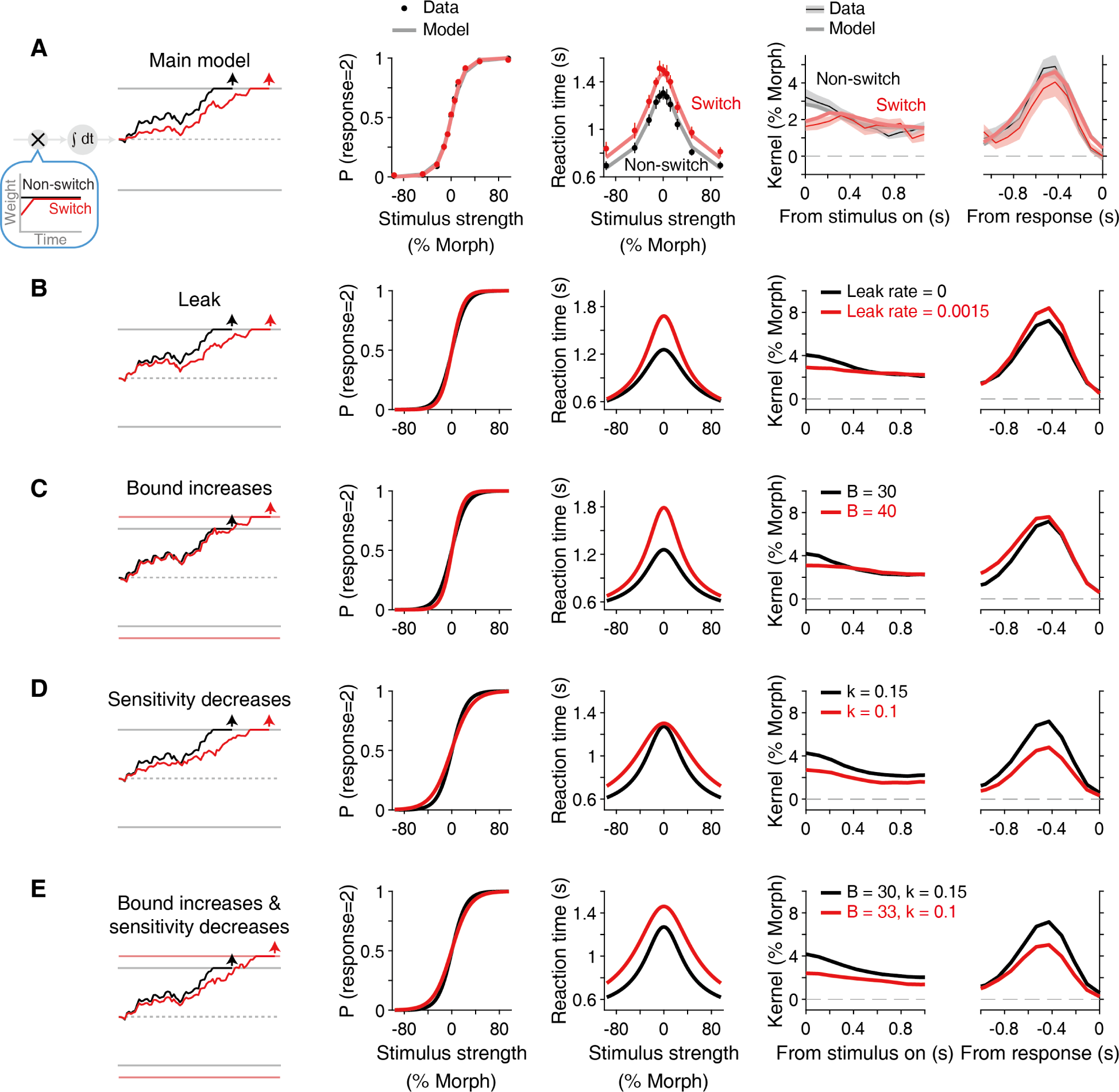
Only the model with initial weight reduction accounts for behavioral data. To test whether any alternative models explain the behavioral data, we varied a number of model parameters (the left column) and plotted the expected changes in psychometric, chronometric functions, and psychophysical kernels (the middle and right columns). Other than our main model, no model produced results that qualitatively matched the actual data. The plots are based on Monte Carlo simulations for demonstration purposes. We fitted each model to data and found that the fitting performance was poorer than our main model. The decrease in log-likelihood for each alternative model was 53.5 (leak; **B**), 35.0 (bound height; **C**), 35.6 (sensitivity; **D**), 8.0 (bound and sensitivity; **E**), indicating that our main model was at least ∼ 3, 000 times more likely than any other models. The default model parameters in the simulations were k = 0.15, B = 30, mean of T_0_ = 400, SD of T_0_ = 133, leak rate = 0. (legend continued on next page) (**A**) Experimental results and the fits of our main model (the same as in Fig. 3). (**B**) Adding an information leak in the evidence accumulation (Eq. 14) delays reaction times (RTs). However, the leak increases RTs especially for difficult stimuli because the model takes more time to accumulate evidence and thus becomes more sensitive to the leak. Furthermore, the amplitudes of the psychophysical kernels are reduced for a longer time window, not limited to the first 200-300 ms. (**C**) Setting the decision bound higher increases RTs, but it particularly affects RTs for difficult stimuli. It also increases choice accuracy. (**D**) Setting the sensitivity lower also increases RTs, but the effect is then more pronounced for easier stimuli. This also decreases choice accuracy. (**E**) Increasing the decision bound and sensitivity balances out these two effects and leads to an increase in RTs for all stimulus strengths while maintaining the choice accuracy. This appears to be consistent with the experimental data, but the kernels of the model show reduced amplitudes throughout the stimulus presentation because individual pieces of evidence have less influence on the model outputs.

**Figure S4:**
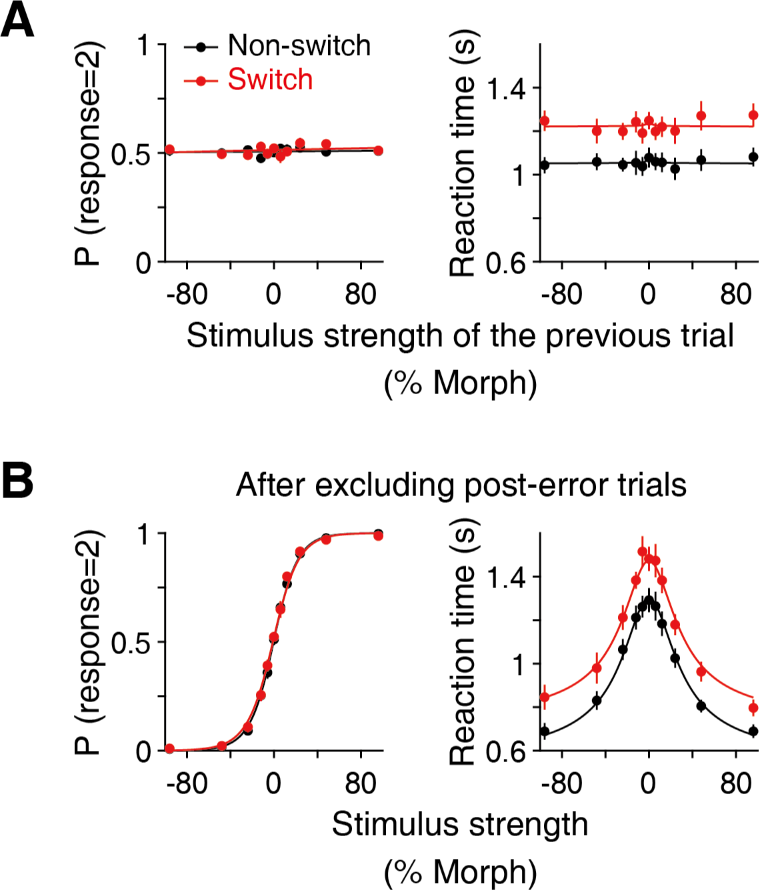
Switch cost depends on neither stimulus history nor past feedback. (**A, B**) If the observed reduction in initial sensory weighting was caused by a stimulus history effect, such as adaptation or priming, the stimulus strength of the previous trial should correlate with the reaction time (RT) increase. For example, previous exposure to a strong stimulus along one stimulus axis could facilitate/hinder decision making along the same/different axis in the next trial. However, we did not observe such an effect in our data. The plots show the data from the main face categorization task (Fig. 1). Error bars indicate S.E.M. across participants. (**C, D**) Past feedback is also known to affect RTs. For example, RTs are often slower after error trials (post-error slowing; Purcell and Kiani 2016). To exclude this effect, we plotted choices and RTs after removing the trials following an error. The result showed no noticeable difference from the main figure (Fig. 2A).

**Figure S5:**
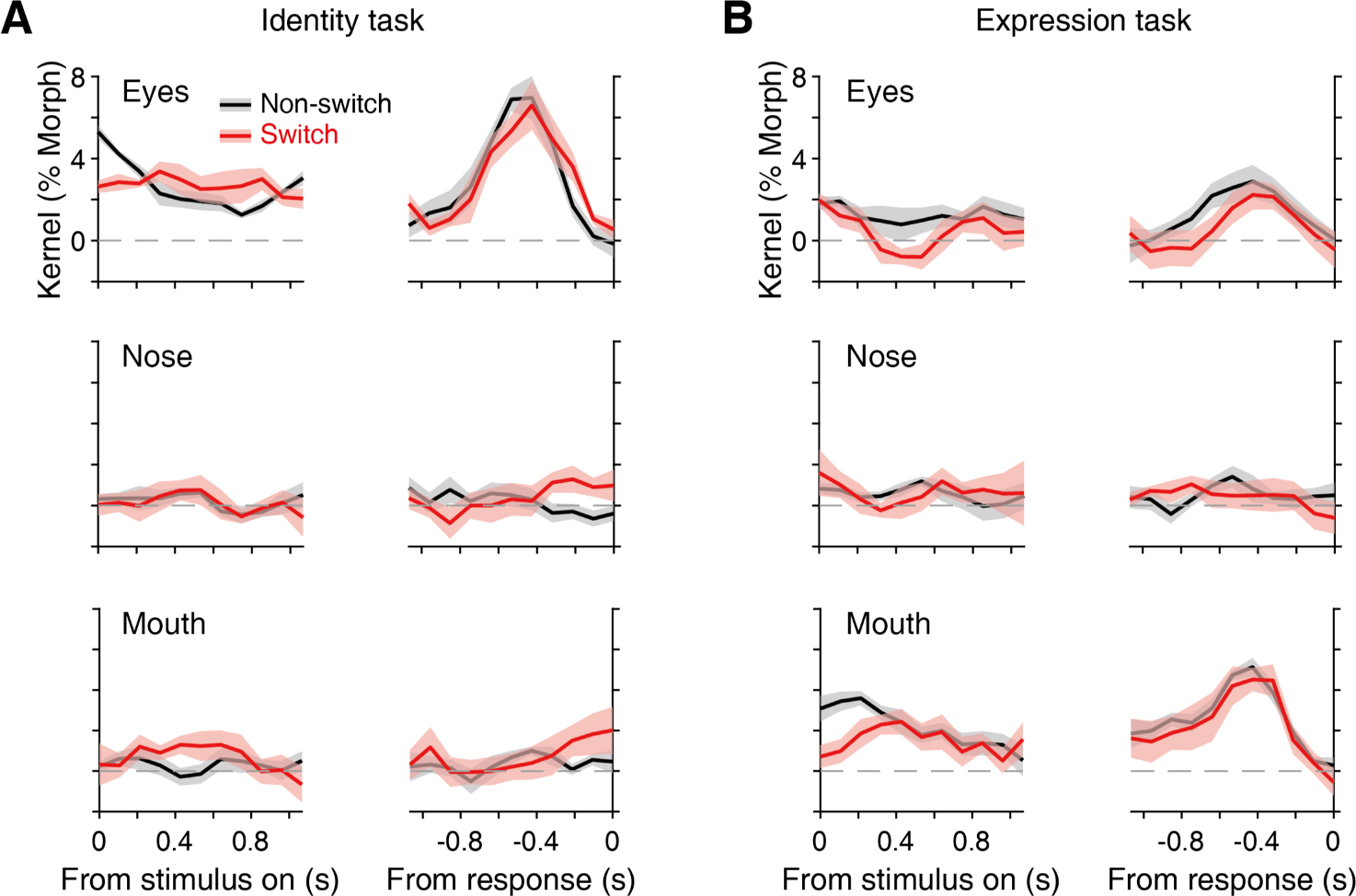
Psychophysical kernels for each task rule and facial feature. (**A, B**) Psychophysical kernels for individual facial features (eyes, nose, and mouth) in the identity (A) and expression (B) categorization rules in the main face task (Fig. 1). Consistent with previous work, participants predominantly weighted the eyes for identity categorization and the mouth for expression categorization (Okazawa et al. 2021b; Schyns et al. 2002) The initial reduction in sensory weight is evident in the kernels for individual features. In the main figure, we averaged the stimulus fluctuations of the features to focus on the time course of the psychophysical kernels (see Methods). Shading indicates S.E.M. across participants.

